# Mutation-Agnostic Base Editing of the Progerin Farnesylation Site Rescues Hutchinson-Gilford Progeria Syndrome Phenotypes in Neuromuscular Organoids

**DOI:** 10.1101/2025.10.16.682736

**Authors:** Dong-Woo Kim, Eun-Ji Kwon, Beom-Jin Jeon, Dong-Hyeok Kwon, Goo Jang, Youngyoon Yoon, Yuna Hwang, Hyukjin Lee, Hyuk-Jin Cha

## Abstract

Hutchinson Gilford progeria syndrome (HGPS) is a fatal premature aging disorder caused by pathogenic farnesylated lamin A variants that disrupt nuclear architecture and DNA repair. Current therapies, including farnesyltransferase inhibitors, provide only modest survival benefits and lack molecular specificity, while mutation-specific genome-editing strategies cannot address atypical laminopathies. Here, we develop Farnesylation Amino acid Targeted Editing (FATE), a mutation-agnostic base-editing platform that selectively disrupts the *LMNA* farnesylation motif without affecting other farnesylated proteins. Using isogenic human pluripotent stem cell derived neuromuscular organoids (NMOs), we reveal muscle-specific progerin accumulation that sequesters 53BP1 and abolishes DNA damage foci formation. FATE eliminates perinuclear progerin, restores 53BP1 mobility, reconstitutes DNA repair foci, and normalizes heterochromatin architecture. Importantly, transient delivery of FATE mRNA conjugated with lipid nanoparticles to HGPS NMOs achieves efficient base editing and phenotypic rescue. These findings establish FATE as a mutation-independent therapeutic strategy targeting a fundamental pathogenic mechanism in HGPS and provide a proof-of-concept for RNA-based in situ genome editing in progeroid disease.

## Introduction

Hutchinson–Gilford progeria syndrome (HGPS; OMIM #176670) is a rare genetic disorder that causes premature aging and leads to early mortality, typically within the first two decades of life ^1^. In over 90% of cases, a de novo LMNA mutation (c.1824C>T; p.G608G) activates a cryptic splice donor in exon 11, producing progerin—a truncated form of lamin A that retains its C-terminal cysteine-serine-isoleucine-methionine (CSIM) ‘CAAX’ motif and hence remains permanently farnesylated ^2,3^. Because progerin lacks the cleavage site for the ZMPSTE24 metalloprotease, it escapes normal post-translational processing and accumulates at the inner nuclear membrane, causing toxic nuclear-envelope deformation ^1^. Beyond the classical c.1824C>T mutation, other LMNA variants (e.g., c.1821G>A, c.1822G>A, c.1868C>G, c.1940T>G, c.1968G>A) are known to impair ZMPSTE24-mediated cleavage, producing farnesylated lamin A variants with progerin-like properties^4^. Likewise, ZMPSTE24 loss-of-function mutations result in accumulation of farnesylated prelamin A that causes related laminopathies such as restrictive dermopathy and mandibuloacral dysplasia ^5,6^. Collectively, despite their distinct genetic causes, all of these conditions share a common pathogenic mechanism: persistent lamin A farnesylation.

HGPS phenotypes are most pronounced in mesoderm-derived tissues such as vascular smooth muscle, bone, skin dermis, and adipose tissue ^7,8^. The persistent accumulation of progerin disrupts the nuclear lamina, causing mechanical fragility, aberrant mitosis, and accumulation of DNA damage ^9^. Impaired recruitment of DNA repair proteins such as 53BP1 and RAD51 compromises double-strand break (DSB) repair and triggers checkpoint activation, ultimately promoting chronic genomic instability and senescence ^9^. Simultaneously, heterochromatin erosion and detachment of lamina-associated domains alter gene expression programs ^10,11^. Together, these nuclear and epigenetic perturbations drive progressive tissue degeneration and functional decline. The cardiovascular system, particularly vascular smooth muscle cells, is especially vulnerable to these stresses, and its degeneration underlies the cardiovascular complications that constitute the leading cause of death in HGPS patients ^7^.

The first and only FDA-approved therapy for HGPS, lonafarnib (Zokinvy), is a farnesyltransferase inhibitor (FTI) that blocks progerin farnesylation, improves nuclear morphology, and extends median survival by approximately 2.5 years ^12,13^. This therapeutic benefit underscores the pathogenicity of farnesylated progerin; however, the activity of lonafarnib is not specific to progerin alone. Long-term FTI use is therefore constrained by systemic toxicity, as farnesylation is essential for the proper function of multiple cellular proteins, including small GTPases (e.g., RAS isoforms and RHOE) and nuclear lamins (LMNB1 and LMNB2). Lonafarnib-induced hypo-farnesylation of these proteins impairs signal transduction, cytoskeletal dynamics, and vesicular trafficking, with consequent off-target effects such as gastrointestinal symptoms, anemia, and myelosuppression ^12,14^. Other therapeutic strategies have emerged, each of which also carries limitations. Rapamycin, which promotes progerin clearance via autophagy, has shown efficacy in preclinical models but also possesses broad immunosuppressive activity that hampers its clinical utility ^15^. Mutation-specific approaches include adenine base editor (ABE) to directly revert the pathogenic c.1824C>T mutation ^16^, which is applicable only to classical HGPS carrying this specific variant. Other strategies with potentially broader applicability include antisense oligonucleotides that block the aberrant splice donor in LMNA ^17,18^, which could in principle be adapted to different splice-activating mutations, and CRISPR–Cas9-mediated selective disruption of lamin A while sparing lamin C ^19^, which is likewise independent of the underlying mutation but carries risks of large genomic deletions or rearrangements^20^. However, none of these approaches directly addresses disorders such as ZMPSTE24 deficiency, which also result in the accumulation of farnesylated prelamin A.

Here, we introduce Farnesylation Amino-acid Targeted Editing (FATE), a base editing approach that targets the CSIM motif of LMNA rather than a specific pathogenic nucleotide. By installing a cysteine-to-arginine substitution in the CAAX box, FATE abolishes farnesylation of not only prelamin A but also progerin, thereby preventing the aberrant retention at the inner nuclear membrane regardless of upstream mutation. Using neuromuscular organoids (NMOs) derived from isogenic LMNA-mutant human pluripotent stem cells (hPSCs), we demonstrate that FATE eliminates perinuclear progerin aggregates and restores DNA damage repair foci. Because FATE disrupts a shared pathogenic post-translational modification rather than a patient-specific LMNA mutation, it provides a mutation-agnostic, potentially pan-laminopathy therapy, expanding the scope of genome editing beyond existing HGPS-targeted approaches.

## Results

### Establishment of two isogenic NMO models with mesoderm-specific progerin expression

To overcome the limited availability of primary fibroblasts from HGPS patients for in vitro studies ^11,15^, human induced pluripotent stem cells (iPSCs) derived from HGPS individuals are widely utilized to generate cellular models for mechanistic investigations ^21,22^. To minimize confounding effects from genetic background variability ^23,24^, we established two isogenic pairs of human pluripotent stem cells (hPSCs) differing only at the LMNA locus:

1. a pair of patient-derived iPSCs carrying the HGPS c.1824C>T mutation and its corrected counterpart (henceforth HGPS- and Edit-iPSCs), and ii) a pair of hESCs harboring either wild-type or engineered c.1824C>T mutant alleles (henceforth WT- and Mutant-hESCs) (Fig. 1a). These isogenic hPSC lines were generated using prime editor as previously described ^25,26^ (Fig. 1b). Consistent with earlier reports ^21^, undifferentiated hPSCs exhibited negligible LMNA expression and thus no detectable progerin production, even in HGPS-iPSCs (Extended Data Fig. 1a).

**Fig. 1:**
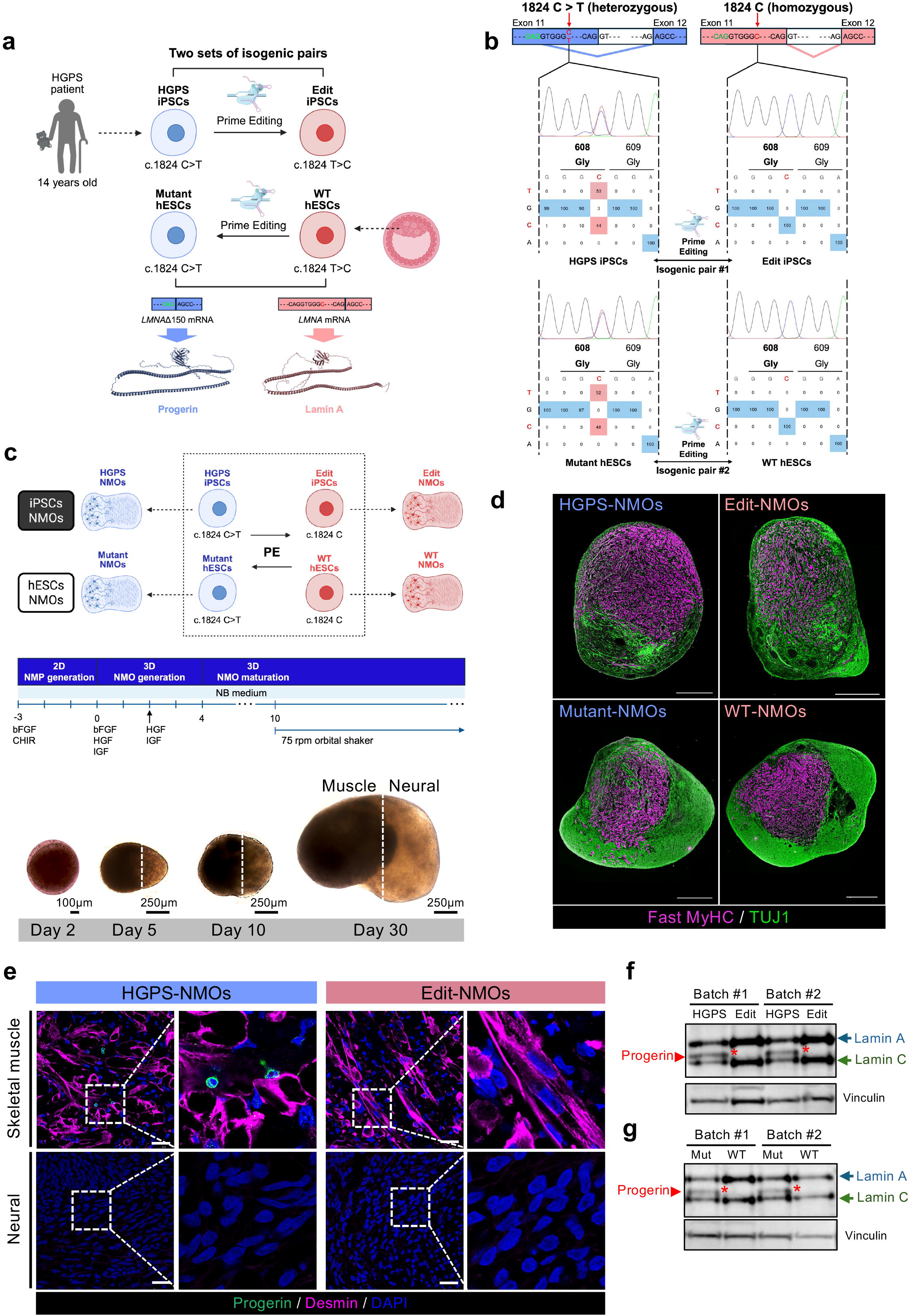
Establishment of two sets of isogenic NMO pairs, with progerin Expression restricted to the muscular compartment of HGPS- and Mutant-NMOs. **a**, Graphical description showing the establishment of two sets of isogenic hPSC pairs. **b**, Sanger sequencing results showing the genotypes of the c.1824 region for each hPSC line. **c**, Graphical overview of neuromuscular organoid (NMO) differentiation. Bright-field images of representative organoid at different stages (days 2, 5, 10 and 30.) **d**, Immunofluorescence image showing established day 70 NMOs derived from two sets of isogenic hPSCs. Magenta indicates fast myosin heavy chain (muscle marker), and green indicates Tuj1 (neuron marker). Scale bar, 500 μm. **e**, Immunofluorescence images of day 40 HGPS and Edit NMOs stained with Progerin (green), Desmin (magenta), and DAPI (blue). Scale bar, 25 μm. **f, g**, Immunoblot analysis of NMOs derived from iPSC and hESC pairs, showing Lamin A/C and Vinculin. Red asterisks indicate progerin-specific bands.

Given that HGPS pathologies are prominent in mesoderm-derived tissues such as skeletal muscle ^7,8^, whereas ectodermal tissues such as neurons remain largely spared, we generated NMOs from the isogenic hPSC lines using established protocols ^27^ (Fig. 1c) to allow the simultaneous modeling of mesodermal (i.e., muscular) and ectodermal (neural) lineages within a shared genetic background. We established two NMO pairs, HGPS- vs. Edit-NMOs and Mutant- vs. WT NMOs, which exhibited comparable compartmentalization of neural and muscular tissues, as confirmed by expression of lineage-specific markers (Fig. 1d and Extended Data Fig. 1b) and ultrastructural features such as M- and Z-bands, synaptic vesicles, and neural filaments (Extended Data Fig. 1c). Despite this similar tissue organization, immunofluorescence revealed progerin accumulation to occur exclusively in the muscular compartments of HGPS-NMOs (Fig. 1e) and Mutant-NMOs (Extended Data Fig. 1d). Immunoblotting further validated robust progerin expression in the disease NMOs (HGPS: Fig. 1f; Mutant: Fig. 1g), whereas none was detected in their respective controls.

### Myopathy-associated gene signatures in the muscle compartment of HGPS-NMOs

Given the selective accumulation of progerin in the muscle compartment of HGPS-NMOs (Fig. 1), we performed single-cell RNA sequencing (scRNA-seq) to further characterize cell-type-specific transcriptional alterations. Graph-based clustering of single-cell transcriptomes identified twelve distinct cell subtypes: skeletal muscle (CL3, CL7), neural cells (CL1, CL2, CL4, CL5), sclerotome (CL0), endothelial (CL9), and various epithelial populations (CL6, CL8, CL10, CL11) (Fig. 2a). The overall distribution of cell types was comparable between HGPS- and Edit-NMOs (Fig. 2b), and lineage-specific marker gene expression confirmed NMO-derived populations to have well-defined differentiation trajectories toward muscle and neural lineages (Fig. 2c). Within the skeletal muscle lineage, we identified a continuum of differentiation states—from myogenic progenitor/satellite-like cells (cluster 1) through intermediate myocytes (cluster 2) to mature contractile skeletal muscle fibers (cluster 3) (Fig. 2d, e). Notably, HGPS-NMOs exhibited a reduced proportion of myogenic progenitors compared to Edit-NMOs, accompanied by increased fractions of differentiated myocytes and mature muscle fibers (Fig. 2d). This shift in cellular composition likely reflects an adaptive regenerative response to chronic muscle damage, and resembles patterns observed in scRNA-seq analyses of Duchenne muscular dystrophy ^28^. Consistent with this interpretation, gene set enrichment analysis revealed significant enrichment of myopathy-associated gene signatures specifically in the muscle lineage of HGPS-NMOs (Fig. 2f); in addition, HGPS muscle populations showed marked upregulation of genes involved in the ‘Striated Muscle Contraction Pathway,’ including actinin family members. These findings align with transcriptomic hallmarks of muscle degeneration and compensatory remodeling.

**Fig. 2:**
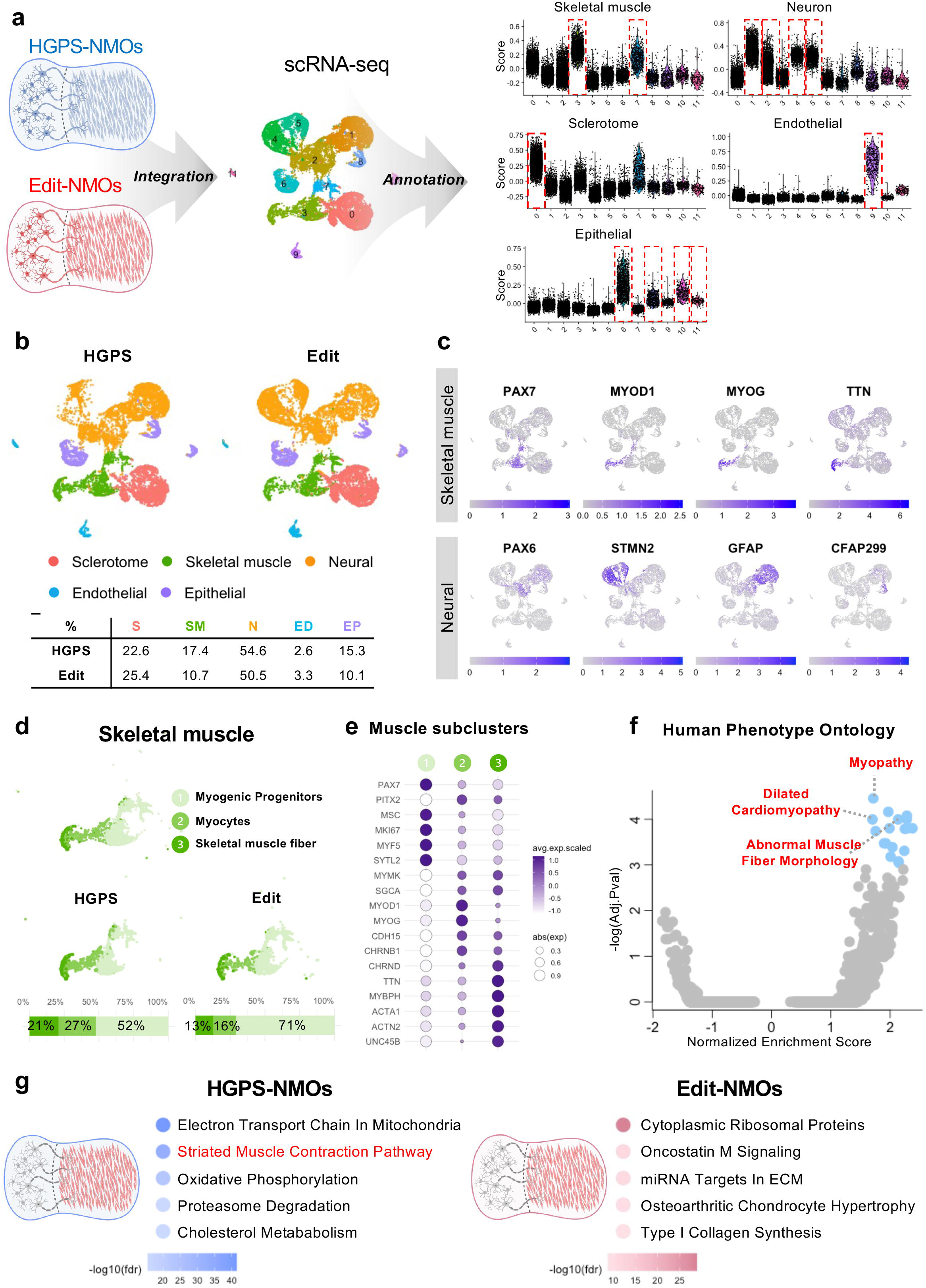
Single-cell analysis of skeletal muscle cellular composition and developmental trajectories in HGPS and Edit NMOs. **a**, Integration, and clustering of single-cell transcriptomes from HGPS- and Edit-NMOs. Cell clusters were annotated based on the enrichment of canonical cell type marker genes. **b**, Comparison of cellular composition between HGPS- and Edit-NMOs. **c**, Identification of skeletal muscle and neural cell populations using lineage-specific markers. **d**, Sub-clustering of the skeletal muscle trajectories reveals 3 distinct subpopulations. Shades of green indicate skeletal muscle subclusters. Relative proportions of each subcluster in HGPS organoids are shown below. **e**, Dot plot showing representative gene expression across three muscle subclusters. **f**, Human Phenotype Ontology terms enriched in HGPS muscle cells compared to Edit controls. **g**, Pathway enrichment analysis of skeletal muscle compartments in HGPS- and Edit-NMOs.

In contrast, the neural lineage—comprising trunk neural crest derivatives (cluster 7) that further differentiated into glial and Schwann cells (cluster 8) (Extended Data Fig. 2a, b)—exhibited no enrichment of disease-related gene sets. Specifically, in HGPS neural populations, both gene enrichment and differential expression analysis failed to reveal transcriptomic signatures associated with pathological phenotypes (Extended Data Fig. 2c, d). Overall, this muscle-specific, neuron-sparing transcriptomic pathology highlights the value of the HGPS-NMO model for dissecting lineage-restricted mechanisms in progeroid disease.

### Impairment of DNA damage repair machinery in the muscle compartment of HGPS-NMOs

Following identification of robust progerin accumulation in the muscle compartment (Fig. 2f) and the muscle-specific HGPS transcriptomic signature (Fig. 3), we investigated the underlying mechanisms by analyzing gene sets with reduced enrichment specifically in HGPS-derived muscle cells compared to Edit controls. Gene sets exclusively altered in the muscle compartment of HGPS—either upregulated in Edit-NMOs or downregulated in HGPS-NMOs—likely represent HGPS-specific transcriptional responses. Notably, gene sets that were upregulated in the muscle compartment of Edit-NMOs compared to HGPS-NMOs, but not in the neural compartment, were enriched for DNA damage repair pathways (highlighted in red, Fig. 3a), consistent with previous reports linking genome instability to HGPS pathogenesis ^9,29^. Additionally, gene sets downregulated in HGPS-NMOs included pathways associated with metabolism and autophagy (highlighted in green, Extended Data Fig. 3a), both of which have been implicated in skeletal muscle dysfunction in HGPS mouse models ^15,30,31^.

**Fig. 3:**
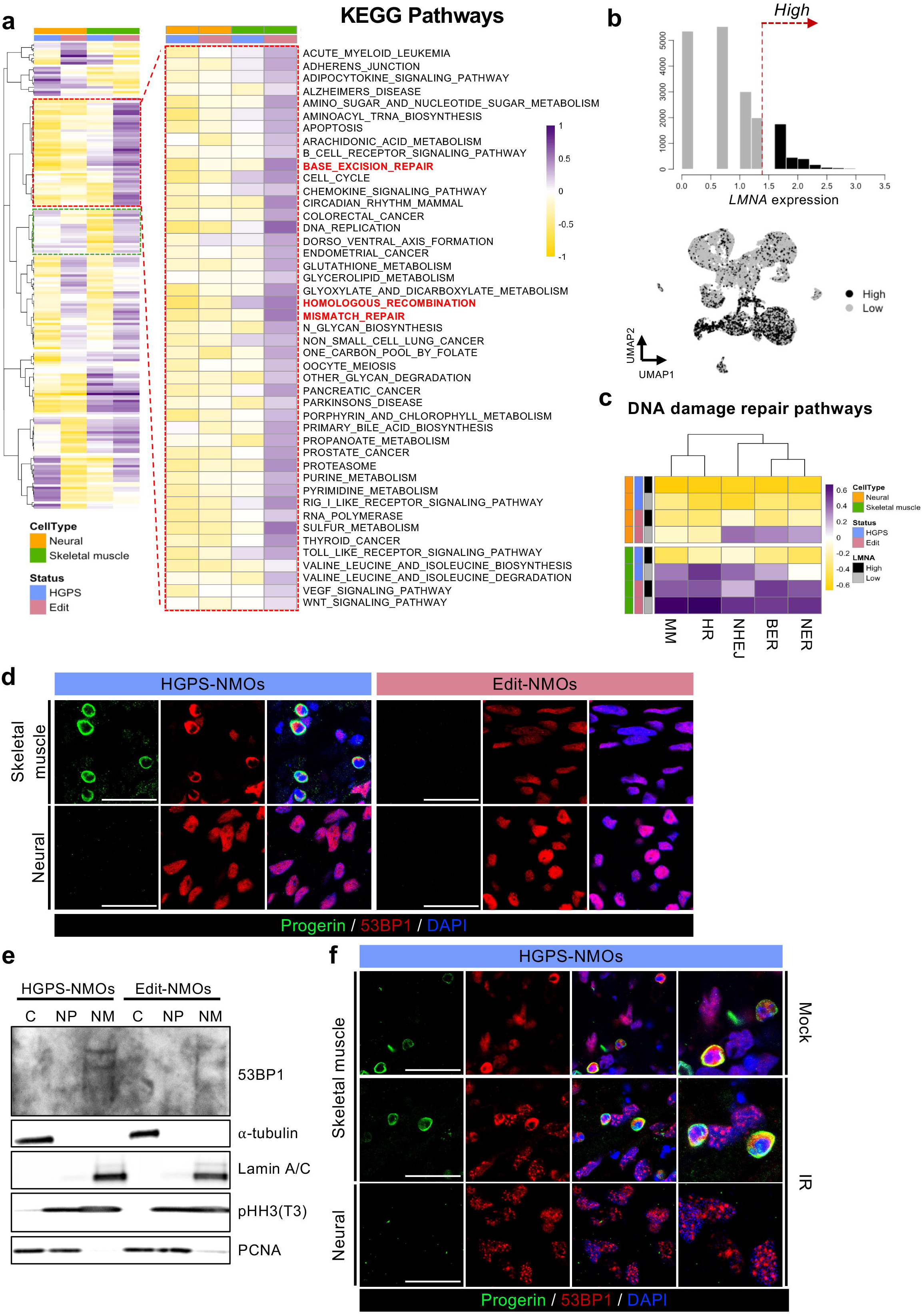
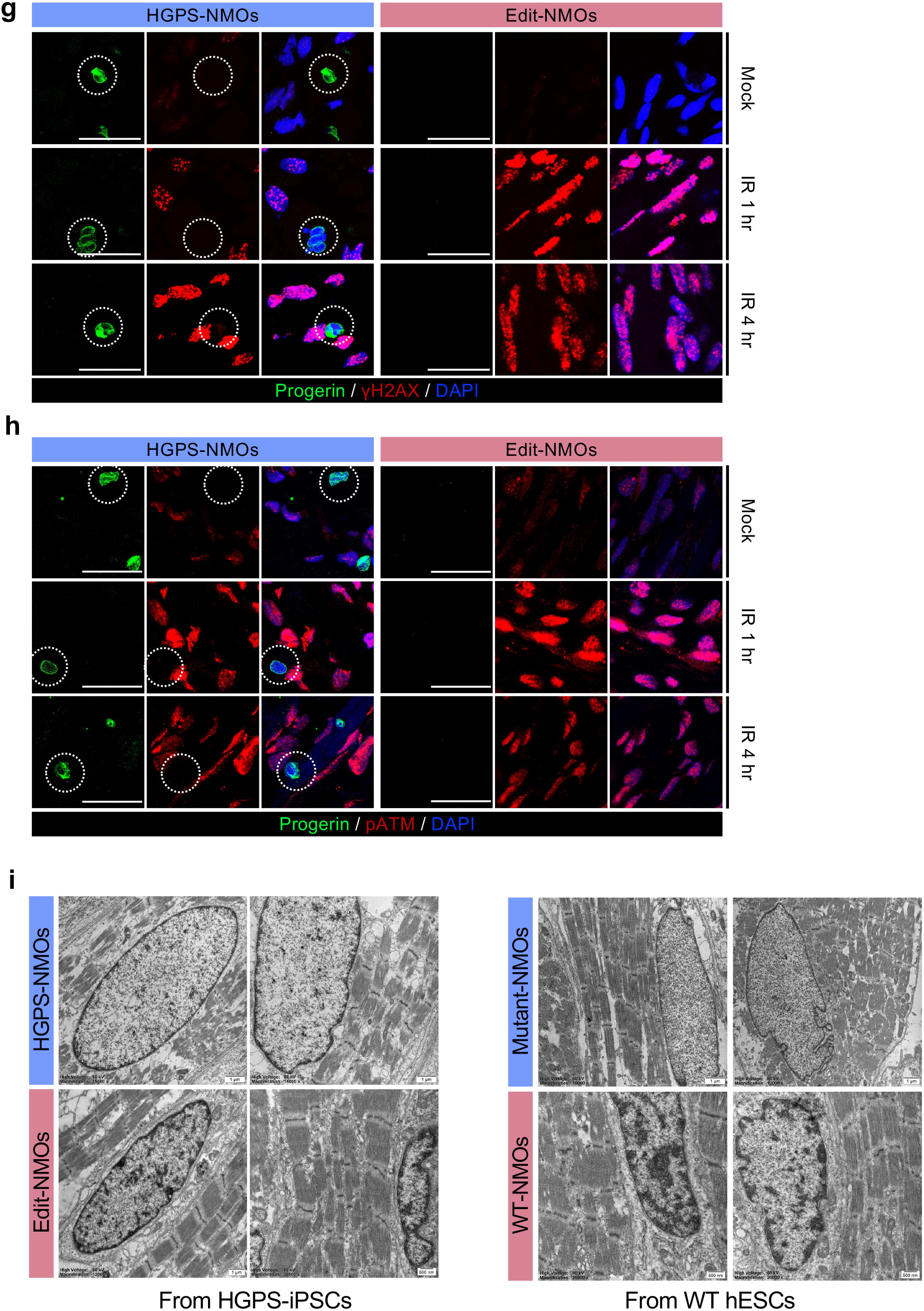
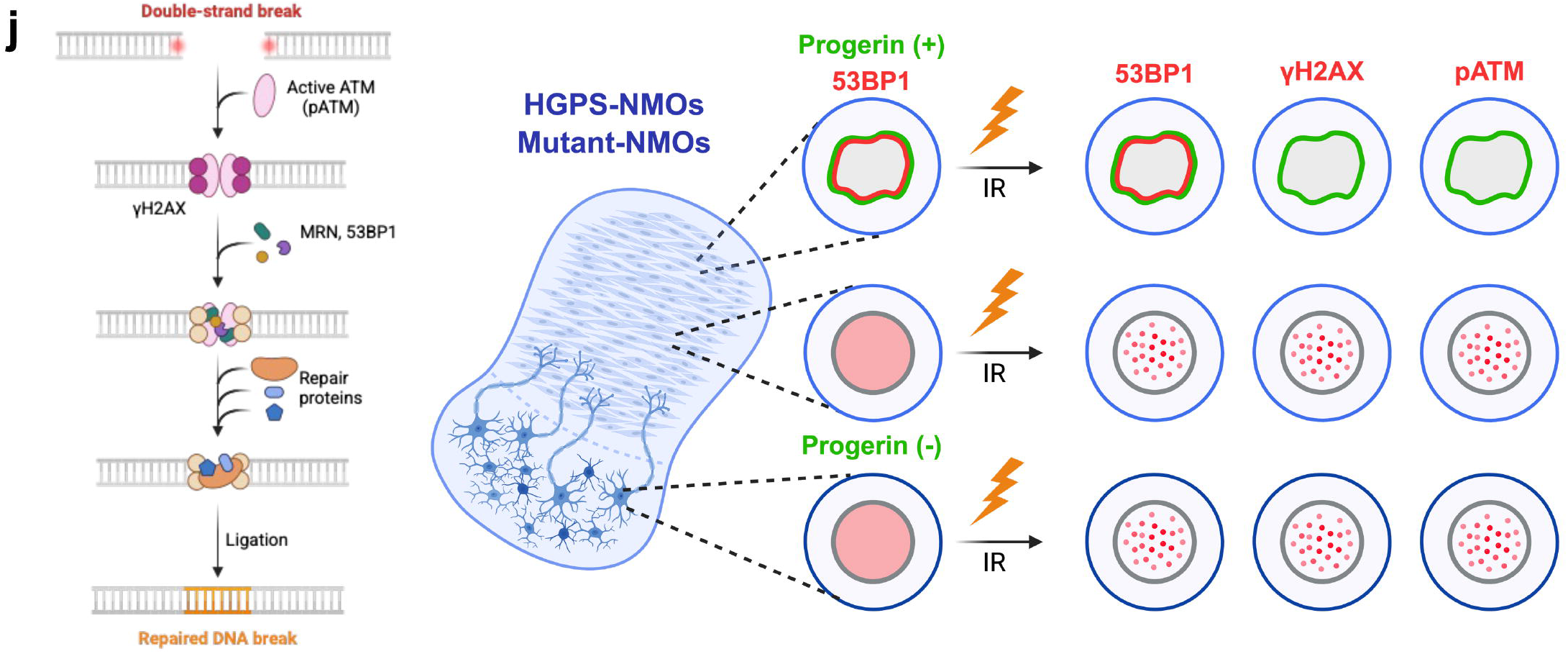
Impaired foci formation of DNA damage repair machinery and heterochromatin loss in the muscle compartment of HGPS-NMOs. **a**, Heatmap showing GSVA-based KEGG pathway enrichment in neural and muscle compartments of HGPS and Edit organoids. Gene sets related to DNA repair pathways are highlighted in red, while those associated with metabolism and autophagy are shown in green. **b**, Cells with *LMNA* expression in the top 25% were designated as the “*LMNA*-high” group (top). UMAP visualization shows the distribution of *LMNA*-high and *LMNA*-low groups across organoids (bottom). **c**, Heatmap displaying enrichment of DNA repair–related gene sets in *LMNA*-high and *LMNA*-low groups within the neural and muscle compartments of HGPS and Edit organoids. **d**, Immunofluorescence images of HGPS and Edit NMOs stained with Progerin (green), 53BP1 (red), and DAPI (blue). Scale bar, 25 μm. **e**, Immunoblot analysis of NMOs derived from iPSCs pairs, showing α-tubulin, Lamin A/C, pHH3 (T3), PCNA, and 53BP1. **f**, Immunofluorescence images of HGPS and Edit NMOs stained with Progerin (green), DAPI (blue), and 53BP1 (red), under mock and 5 Gy X-ray irradiation for 1 hour. Scale bar, 25 μm. **g**, **h**, Immunofluorescence images of HGPS and Edit NMOs stained with Progerin (green), DAPI (blue), and each of the following DNA damage markers (red) under mock, 1-hour, and 4-hour conditions following 5 Gy X-ray irradiation: (**g**) γH2AX; (**h**) phosphorylated ATM (pATM, S1981). Scale bar, 25 μm. **i**, Transmission electron microscopy (TEM) images showing the nuclear heterochromatin structure in the muscle region of neuromuscular organoids (NMOs) derived from two isogenic HGPS hPSC pairs. **j**, Graphical description illustrating defective localization of DNA damage markers 53BP1, γH2AX, and phosphorylated ATM (pATM, S1981) in HGPS-NMOs and mutant-NMOs expressing progerin, under both mock and 5 Gy X-ray irradiation conditions, indicating impaired DNA double-strand break response compared to isogenic controls.

Given that HGPS-iPSCs are heterozygous and thus express both wild-type and mutant LMNA alleles at equivalent levels, we hypothesized that for individual cells, progerin expression would scale proportionally with LMNA transcript abundance (Extended Data Fig. 3b). We therefore stratified cells based on LMNA expression, defining the top 25% as “LMNA-high” and the remainder as “LMNA-low” (Fig. 3b). Strikingly, within the muscle compartment of HGPS-NMOs, DNA damage repair gene sets were more suppressed in the LMNA-high group (Fig. 3c), suggesting that elevated LMNA expression—and thus greater progerin burden—may directly contribute to the impaired DNA repair capacity of HGPS muscle cells. This transcriptomic signature was not associated with altered expression of the progerin-regulating splicing factors *SRSF1* and *SRSF2* ^32,33^, which were also not preferentially expressed in the muscle compartment (Extended Data Fig. 3c).

Lamin A and C are known to interact with p53-binding protein 1 (53BP1) ^34^, which is a key mediator of DSB repair through its recruitment to DNA damage foci ^35^; notably, that recruitment can be disrupted by the presence of lamin B ^36^ or prelamin A ^37^. To assess whether progerin impairs 53BP1 localization, we examined 53BP1 distribution in HGPS-NMOs. Intriguingly, in the muscle compartment of HGPS-iPSCs, 53BP1 was selectively sequestered at the nuclear membrane, the same area where progerin was prominently expressed, suggesting its physical displacement from DNA damage sites (Fig. 3d and Extended Data Fig. 3d). Immunoblotting of the nuclear membrane fraction further highlighted this accumulation (Fig. 3e). This persistent sequestration of 53BP1 and its coincidence with progerin accumulation suggests that in HGPS-NMOs, 53BP1 is unable to form DNA damage foci even in response to DSBs. As anticipated, in muscle cells with perinuclear progerin expression (progerin-positive muscle cells) failed to form DNA damage foci after ionizing radiation (IR), with 53BP1 remaining sequestered at the nuclear periphery (Fig. 3f). In contrast, HGPS-NMO neural cells lacking progerin exhibited robust nuclear 53BP1 foci under the same conditions (Fig. 3f). Similarly, γH2AX—a canonical marker of DSBs characterized by phosphorylation at serine 139 ^38^—was absent from the nuclei of progerin-positive muscle cells of HGPS-NMOs (Fig. 3g), but not of cells in the neural compartment (Extended Data Fig. 4a). Likewise, phosphorylated ATM (pATM), the upstream kinase required for γH2AX activation, forming DNA damage foci at Edit-NMOs upon IR, was undetectable in progerin-positive muscle cells of HGPS-NMOs (Fig. 3h). The absence of γH2AX and pATM signals in muscle cells of HGPS-NMOs is attributed to perinuclear retention of ATM alongside progerin even after IR (Extended Data Fig. 4b). Mutant-NMOs exhibited results consistent with HGPS-NMOs: γH2AX and pATM foci were absent in progerin-positive muscle cells (Extended Data Fig. 4c, 4e) but preserved in the neural compartment (Extended Data Fig. 4d). Loss of peripheral heterochromatin architecture—a hallmark of genomic instability in HGPS ^9^—was evident in muscle cells of HGPS- and Mutant-NMOs, but not in WT- or Edit-NMOs (Fig. 3i), and was consistently absent in neurons (Extended Data Fig. 5a). Given the essential roles of ATM-mediated γH2AX formation and 53BP1 recruitment in DSB repair (Fig. 3j, left), persistent sequestration of these factors by progerin likely underlies the chronic DNA damage and genome instability characteristic of HGPS (Fig. 3j, right).

### Specific Farnesylation Amino acid Targeted Editing (FATE) of *LMNA*

Previous gene-editing strategies for HGPS include: i) Cas9-mediated disruption of lamin A, which can eliminate progerin expression but carries the risk of introducing large on-site deletions ^39,40^ or even chromosomal rearrangements ^41^; and ii) ABE, which precisely corrects the pathogenic c.1824C>T; p.G608G mutation but is applicable only to HGPS patients carrying this classical HGPS variant. For patients harboring non-classical HGPS mutations that also result in aberrant progerin production, gene correction requires customized editing strategies—such as near-PAMless ABEs ^42^, prime editor (PE), or cytosine-to-guanine base editors (CGBE)—each of which presents distinct technical challenges including the presence of bystander adenines (Extended Data Fig. 6a). To overcome these limitations and establish a mutation-agnostic approach, we targeted progerin farnesylation—a well-established driver of HGPS pathology—by designing a base editing strategy to disrupt the conserved CSIM motif in LMNA. As proof of principle, we tested the effect of unfarnesylated progerin generated by substituting the cysteine residue with serine in a GFP–progerin reporter cell line. This substitution reduced nuclear membrane retention of progerin (Extended Data Fig. 6b) and restored DNA damage repair foci (Extended Data Fig. 6c), confirming that disrupting the farnesylation motif is a viable alternative to mutation-specific correction. Building on this rationale, we engineered a single-guide RNA (sgRNA) to introduce the arginine (CGC) substitution at position 661 (c.1981T>C; p.C661R) coding cysteine (TGC), thereby preventing farnesylation and abolishing progerin accumulation at the nuclear envelope—a mutation-agnostic strategy we term Farnesylation Amino acid Targeted Editing (FATE) (Fig. 4a, left). This approach circumvents both the genotoxic risks associated with Cas9-induced large deletions and the complexity of tailoring editing tools to individual HGPS variants, while abolishing post-translational farnesylation of progerin and preventing its aberrant retention at the inner nuclear membrane regardless of the upstream mutation (Fig. 4a, right). To identify optimal editing conditions, we evaluated multiple ABE-compatible guide RNAs that positioned the target TGC codon within the editing window of various protospacer adjacent motif (PAM) sequences and ABE versions (Extended Data Fig. 6d). Among these, only the guide RNA sequence TGCTGCAGTTCTGGGGGCTC, which employs a TAA PAM, exhibited significant editing activity in HGPS-iPSCs (c.1824C>T; p.G608G) when paired with SpRY-ABE8e ^43^, and therefore was used for the generation of FATE-iPSCs (c.1824C>T; p.G608G and c.1981T>C; p.C661R).

**Fig. 4:**
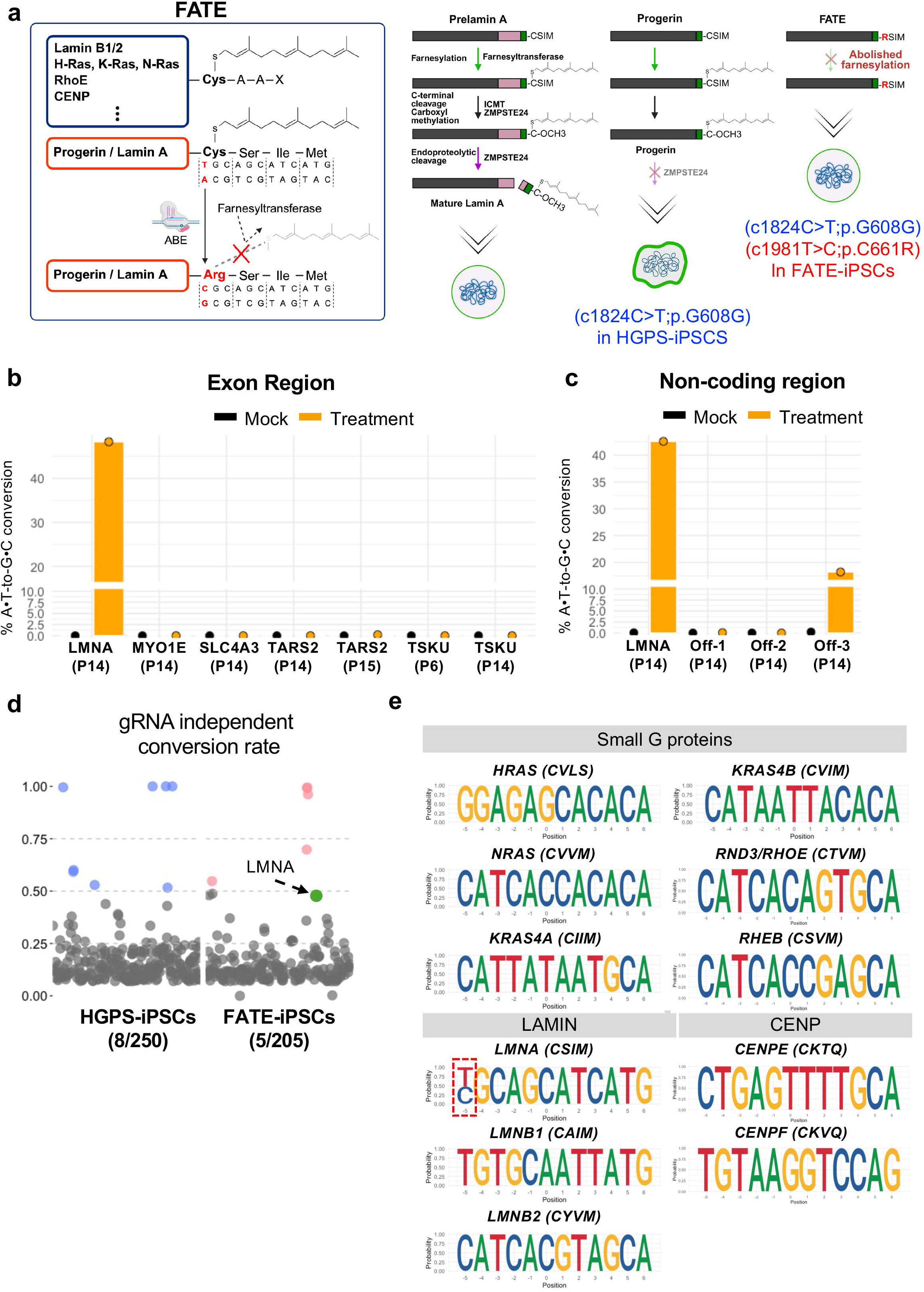
Farnesylation Amino acid Targeted Editing (FATE) exclusively in *LMNA*. **a**, Graphical description showing the Progerin-CAGE strategy, which selectively abolishes farnesylation of progerin by introducing an additional c.1981T>C (p.C661R) mutation into the *LMNA* gene using base editing, without affecting other CAAX box–harboring proteins. **b**, Genome-wide A-to-G conversion frequency at potential gRNA-targetable sites. **c**, A-to-G conversion frequency in non-coding regions of three candidates by targeted deep sequencing. **d**, Dot plot showing genes with high A-to-G conversion rates in HGPS wild-type and HGPS plasmid-treated samples. **e**, DNA sequence logo representation of the CAAX motif in Small GTPase, Lamin, and CENP family genes.

Given that systemic hypo-farnesylation is a major limitation of long-term FTI therapy in HGPS, we first assessed the target selectivity of FATE by whole-exome sequencing (WES) and targeted deep sequencing of HGPS-iPSCs with or without FATE. FATE achieved 48% A-to-G conversion at the intended LMNA site with no guide RNA– dependent off-target edits detected above 0.001% (Fig. 4b and Extended Data Fig. 7a). At sequencing depths >200 reads, WES identified eight and five A-to-G conversion sites in HGPS-iPSCs with or without FATE, respectively; all showed low homology to the guide RNA (maximum 9/20 matching nucleotides) and corresponded to known non-pathogenic SNPs (Fig. 4d and Extended Data Fig. 7c). To capture potential off-target activity in non-coding regions not covered by WES, we performed targeted deep sequencing of three candidate loci. Only one site (Off-3), containing a single mismatch to the guide RNA, exhibited detectable editing (∼18% A•T-to-G•C conversion), and this site resides in an intergenic region, suggesting minimal clinical relevance (Fig. 4c and Extended Data Fig. 7b). Importantly, consistent with its sequence-specific design, FATE did not alter the CAAX motifs of other farnesylated proteins—including small GTPases, lamins, and CENP family members—thereby avoiding the broad suppression of protein farnesylation observed with FTI treatment (Fig. 4e).

### Loss of progerin accumulation and restoration of DNA damage foci in FATE-NMOs

FATE-iPSCs generated from HGPS-iPSCs (Fig. 5a) were further differentiated into FATE-NMOs, which displayed typical skeletal muscle and neural structures (Extended Data Fig. 8a). With these NMOs, we assessed whether disruption of the LMNA farnesylation site could reverse the pathogenic phenotypes observed in HGPS-NMOs, including perinuclear progerin accumulation and the associated failure to form DNA damage foci (Fig. 5b). As anticipated, the characteristic perinuclear progerin aggregation observed in HGPS-NMOs was absent in FATE-NMOs; instead, progerin exhibited a diffuse nuclear localization indicating effective disruption of membrane tethering (Fig. 5c). Notably, this relocalization occurred despite the continued progerin expression in FATE-NMOs, which is distinct from the case of Edit-NMOs, wherein progerin production was eliminated through correction of the pathogenic LMNA mutation (Fig. 5d). Intriguingly, lamin A in FATE-NMOs exhibited an upward shift in electrophoretic mobility relative to both HGPS and Edit-NMOs. This shift likely reflects impaired ZMPSTE24-mediated cleavage, which requires prior farnesylation for substrate recognition and processing ^44^ (Fig. 5d, bottom). Finally, the perinuclear sequestration of 53BP1 observed in the muscle compartment of HGPS-NMOs (Fig. 3d) was abolished in FATE-NMOs (Extended Data Fig. 8b). This was accompanied by reformation of DNA damage foci, indicated by the induction of γH2AX (Fig. 5e) and pATM foci (Fig. 5f) following DSB stimulation, which demonstrates the recovery of DNA repair capacity upon disruption of progerin farnesylation. We applied three-dimensional confocal imaging to visualize the absence or dissociation of progerin and the reconstitution of DNA damage foci before and after IR (Fig. 5g and Video 1-6). TEM further revealed restoration of peripheral heterochromatin structure in the muscle compartment of FATE-NMOs (Extended Data Fig. 8c), a feature severely compromised in HGPS-NMOs and indicative of genomic instability ^9^ (Fig. 5h). Collectively, these results indicate that FATE not only reinstates DNA repair functionality but also promotes broader recovery of nuclear architecture in a lineage-specific manner.

**Fig. 5:**
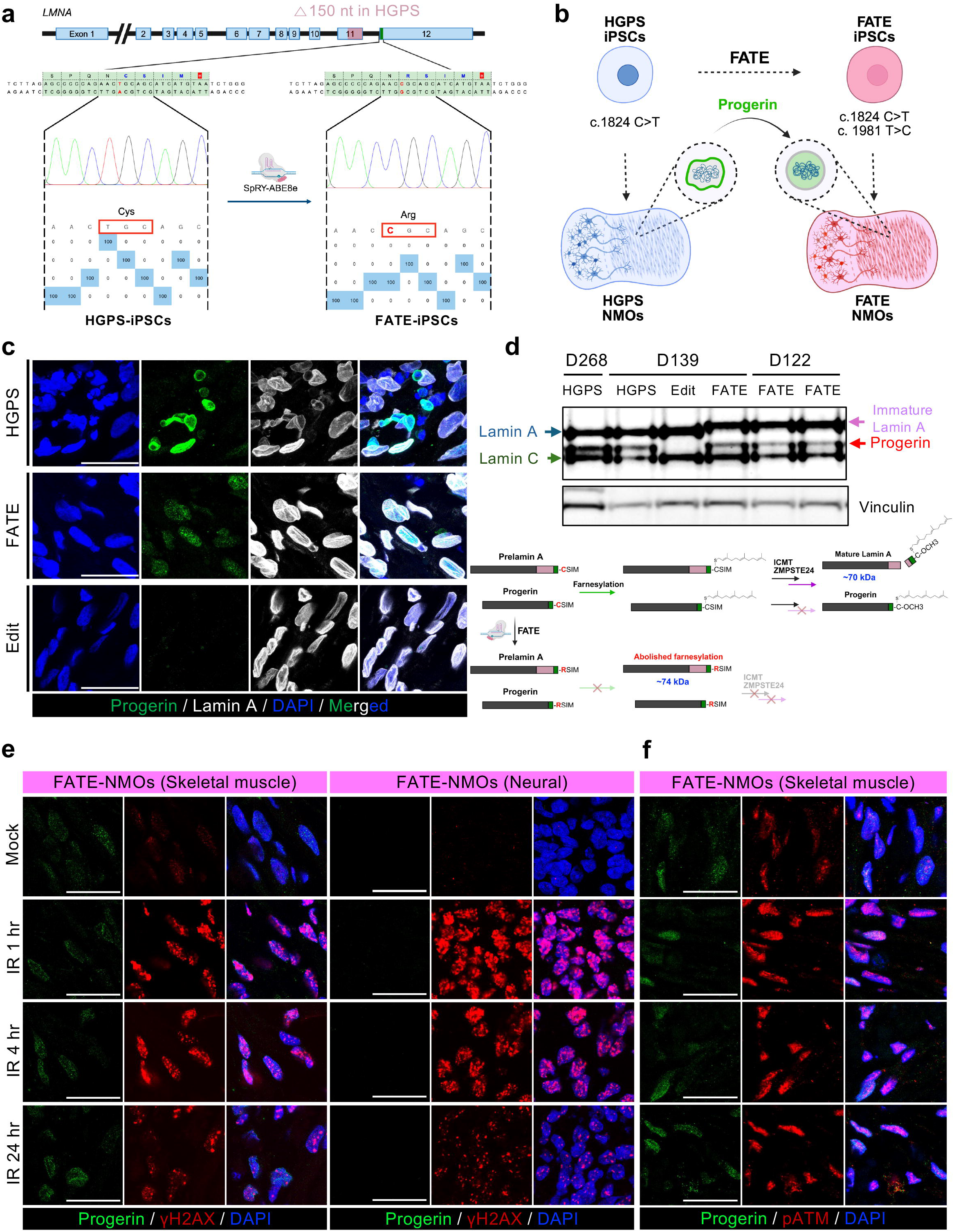
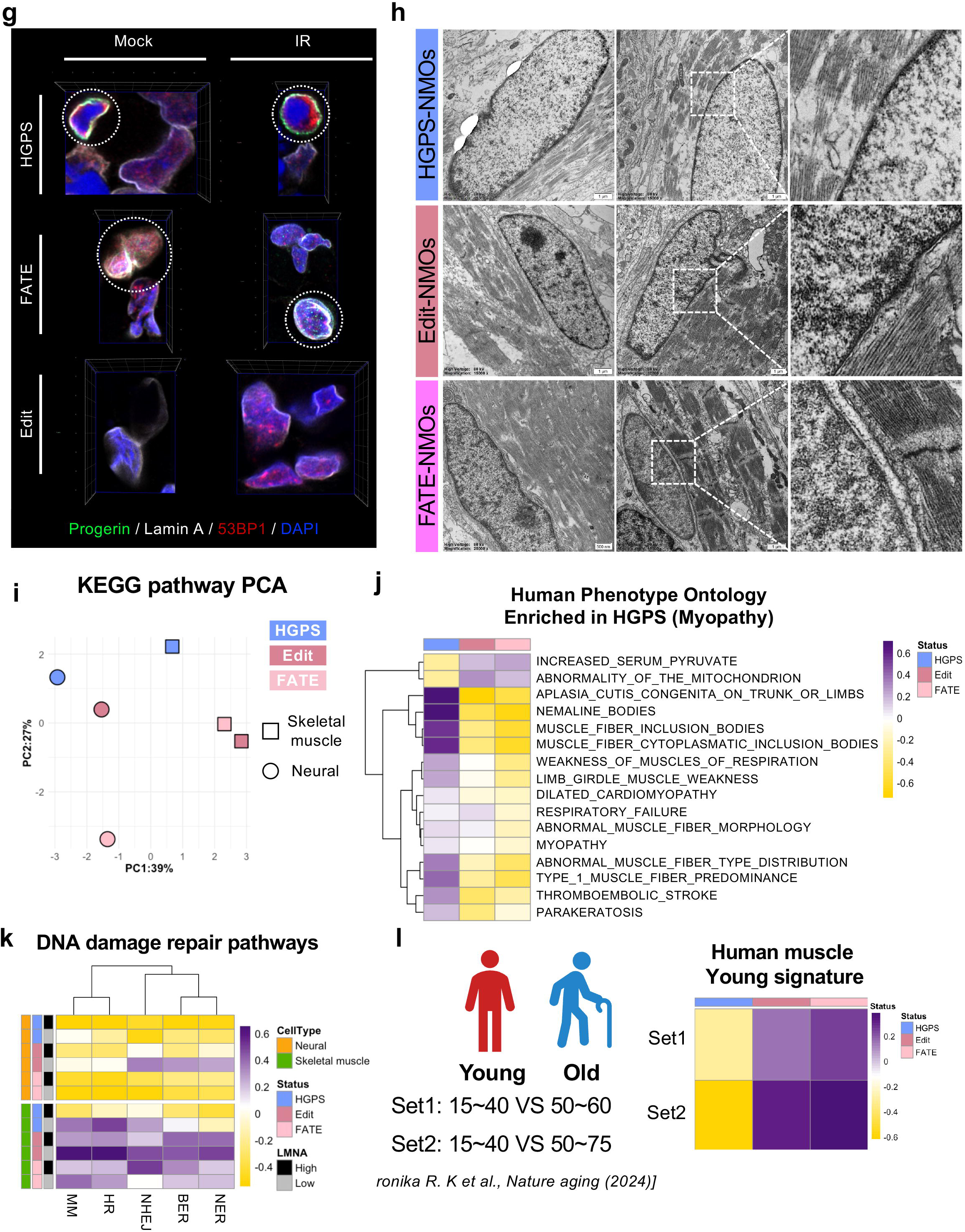
Loss of progerin accumulation, restoration of DNA repair machinery foci formation, recovery of heterochromatin, and normalization of transcriptomic profiles in FATE-NMOs. **a**, Sanger sequencing results showing the establishment of FATE-iPSCs harboring the c.1961T>C (p.C661R) alteration. **b**, Graphical summary showing the establishment of HGPS-NMOs and FATE-NMOs derived from HGPS-iPSCs and FATE-iPSCs, respectively, highlighting the alteration in progerin localization from aberrant nuclear envelope accumulation in HGPS-NMOs to a dispersed nuclear pattern in FATE-NMOs with abolished farnesylation. **c**, Immunofluorescence images of HGPS, Edit, and FATE NMOs stained with Progerin (green), Lamin A (gray), and DAPI (blue). Scale bar, 25 μm. **d**, Immunoblot analysis of HGPS, Edit, and FATE-NMOs showing Lamin A/C and Vinculin (top), and graphical description illustrating changes in post-translational modifications of Lamin A and Progerin induced by FATE (bottom). **e-g**, Immunofluorescence images of CAGE-NMOs stained with Progerin (green), DAPI (blue), and each of the following DNA damage markers (red) under mock, 1-hour, 4-hour, and 24-hour of 5 Gy X-ray irradiation: (**e**) γH2AX, (**f**) phosphorylated ATM (pATM, S1981), and (**g**) 53BP1. **h**, Transmission electron microscopy (TEM) images showing the nuclear heterochromatin structure in the muscle region of neuromuscular organoids (NMOs) derived from two isogenic HGPS hPSC pairs, including FATE-NMOs. **i**, Principal component analysis (PCA) based on GSVA of KEGG pathway enrichment score across HGPS, Edit, and FATE organoid samples. **j**, Heatmap displaying the expression of myopathy-related gene sets across HGPS, gene-edited (Edit), and FATE - treated organoids**. k**, Heatmap displaying enrichment of DNA repair–related gene sets in *LMNA*-high and *LMNA*-low groups within the neural and muscle compartments of HGPS, Edit and FATE organoids. **l**, Heatmap showing enrichment of a “Young” gene expression signature—derived from transcriptomic profiles of young versus old human skeletal muscle—in HGPS, Edit, and FATE organoids.

To systematically characterize the transcriptional changes underlying phenotypic rescue, we performed scRNA-seq on FATE-NMOs and compared them with HGPS- and Edit-NMOs. FATE-NMOs maintained five major cell types, confirming preservation of lineage identities (Extended Data Fig. 8d), and exhibited a continuous differentiation trajectory within both the skeletal muscle (Extended Data Fig. 8e) and neural lineages (Extended Data Fig. 8f), recapitulating normal developmental progression. Notably, the skeletal muscle lineage of FATE-NMOs also showed increased representation of myogenic progenitor populations, mirroring the distribution patterns seen in Edit-NMOs (Fig. 2d).

Principal component analysis of KEGG pathway enrichment profiles indicated the muscle transcriptome of FATE-NMOs to cluster closely with that of Edit-NMOs and diverge from the disease-associated profile of HGPS-NMOs (Fig. 5i). In addition to the reduced myogenic progenitor population observed in HGPS-NMOs (Fig. 2d), the enrichment of myopathy-associated gene signatures (Fig. 2f) and downregulation of DNA damage repair pathways (Fig. 3c) were all restored in FATE-NMOs (Fig. 5j, k). Finally, based on prior evidence that molecular aging accelerates after ages 40 and 60, we classified transcriptomic signatures from bulk RNA-seq of human muscle stem cells ^45^ as either “young” (15–40) or “old” (≥60). Gene set enrichment analysis revealed both FATE-NMOs and Edit-NMOs to have significantly higher enrichment for the “young” signature compared to HGPS-NMOs, indicating rejuvenation of the muscle transcriptome following FATE (Fig. 5l).

### Recovery of HGPS-NMOs by delivery of FATE mRNA

Given the known risks of adeno-associated virus (AAV)–mediated delivery— including potential immunotoxicity, particularly in pediatric patients ^46^, and its limited cargo capacity—we opted to avoid AAV for delivering ABE components. Instead, we tested synthetic mRNA for transient SpRY-ABE8e expression, a platform recently optimized for in vivo gene editing ^47,48^, in the context of FATE in HGPS-iPSCs. For proof of concept, mRNA-based FATE (mRNA-FATE) achieved higher editing efficiency in HGPS-iPSCs compared to plasmid-based delivery (Fig. 6a). We next encapsulated green fluorescent protein (GFP) mRNA-FATE in lipid nanoparticles (LNPs) ^49^, and applied them directly to Day 25 HGPS-NMOs by simple incubation (Fig. 6b). Interestingly, in our neuromuscular organoid model, GFP mRNA-LNPs showed preferential uptake in the muscle compartment. As a result, editing efficiency with ABE8e mRNA-LNP, measured by next-generation sequencing of whole organoids appeared low, reflecting dilution by unedited neuronal and non-muscle cells (Fig. 6c). To address this limitation, we delivered mRNA-LNPs by microinjection directly into the muscle region of NMOs (Fig. 6d), mimicking a local intramuscular injection strategy. To determine an appropriate dosage, GFP mRNA-LNPs were microinjected at 257.58 μg/ml, delivering a total of 515.16 ng (2 μl) of encapsulated mRNA. This resulted in robust GFP expression, confirming the feasibility of localized delivery into organoids (Fig. 6e, left). Following microinjection of ABE8e mRNA-LNPs into Day 26 HGPS-NMOs and an additional 10 days of culture, next-generation sequencing confirmed successful base editing at the target site, with a maximum efficiency of 8.55% (Fig. 6e, right). Given the <10% editing efficiency achieved with FATE mRNA, the observed phenotypic rescue—characterized by loss of perinuclear progerin aggregation and restoration of DNA damage foci—likely reflects a mixed population of edited and unedited cells within the organoids (Fig. 6f). Consistent with genetically edited FATE-NMOs, transient FATE mRNA delivery abolished progerin accumulation at the nuclear membrane, restored its diffuse nuclear distribution, and reestablished DNA damage foci marked by γH2AX (Fig. 6g) and 53BP1 (Fig. 6h). These results demonstrate that even partial, transient mRNA-mediated base editing is sufficient to restore functional DNA repair. Collectively, our findings show that transient FATE mRNA delivery can phenotypically and molecularly rescue HGPS-NMOs, underscoring its translational promise as a mutation-agnostic therapeutic strategy.

**Fig. 6:**
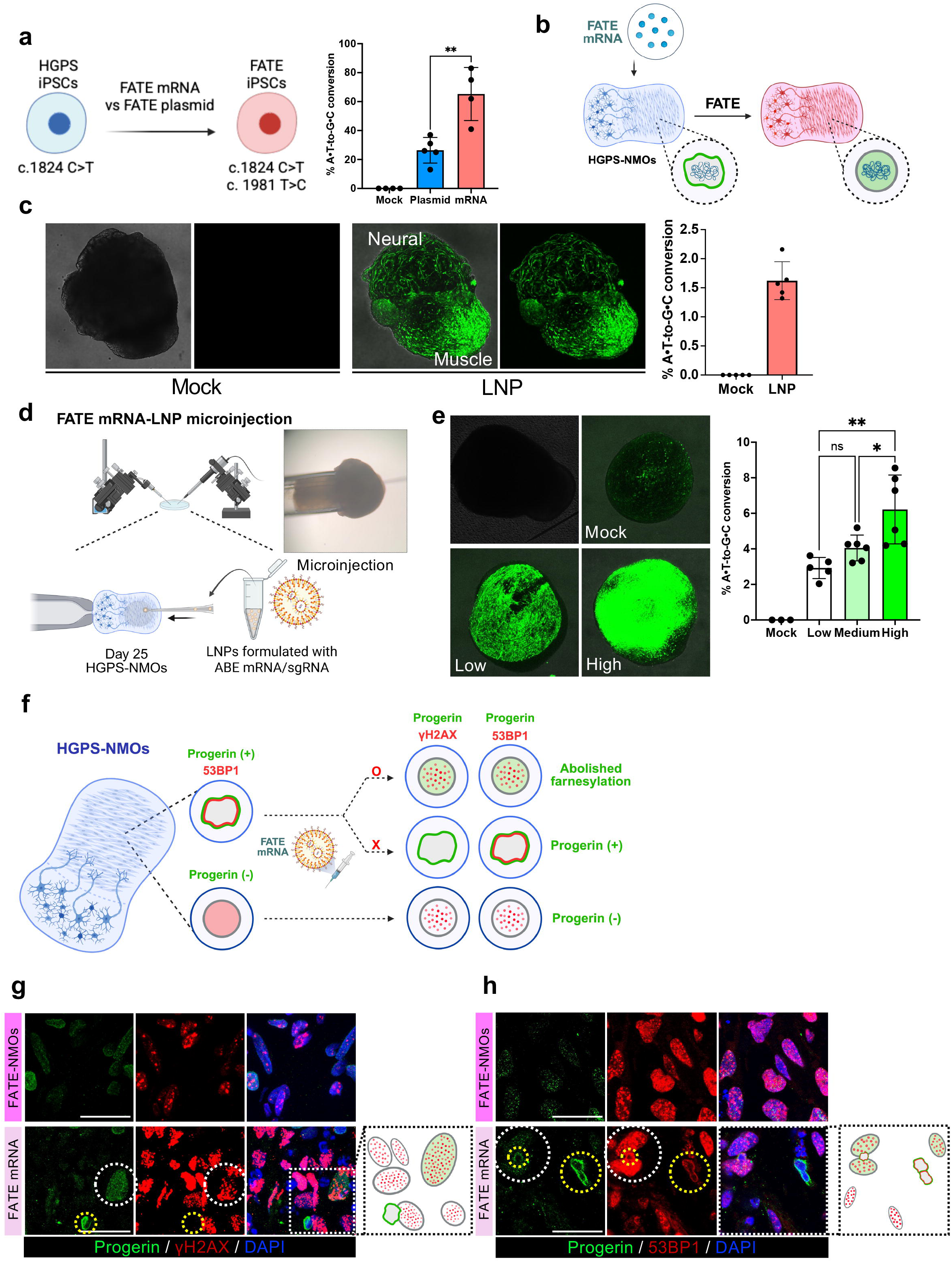
Recovery of disease phenotypes in HGPS-NMOs by delivery of FATE mRNA via direct microinjection. **a**, Graphical description of FATE delivery into HGPS iPSCs via plasmid and mRNA approaches (Fig. 6A, left) and A-to-G conversion rates measured by Sanger sequencing (Fig. 6A, right). **b**, Graphical description illustrating the delivery of FATE mRNA-LNPs into HGPS-NMO. **c**, Confocal images of Day 15 HGPS-NMOs in mock and GFP mRNA–LNP– treated groups, with LNPs diluted in media (Fig. 6c, left), and A-to-G conversion rates measured by NGS after delivery of FATE mRNA–LNPs diluted in media to Day 15 HGPS-NMOs (Fig. 6c, right). **d**, Graphical description of FATE mRNA-LNP Microinjection to HGPS NMOs and bright field image of microinjection. **e**, Confocal images of Day 25 HGPS-NMOs in mock and GFP mRNA–LNP–treated groups following LNP microinjection (Fig. 6e, left), and A-to-G conversion rates measured by NGS 10 days after microinjection of FATE mRNA–LNPs into Day 25 HGPS-NMOs (Fig. 6e, right). (Low: 285.29μg/ml; Medium: 570.58μg/ml; High : 1587.65μg/ml.) **f**, Graphical description of Progerin, γH2AX, and 53BP1 localization in Day 25 HGPS-NMOs after microinjection, comparing edited and non-edited conditions, and observed at Day 70. **g**, **h** Immunofluorescence images of HGPS NMOs after microinjection of FATE mRNA–LNPs at Day 25 and observation at Day 70, stained with Progerin (green), Lamin A (gray), and DAPI (blue), and each of the following DNA damage markers (red) under 1 hour of 5 Gy X-ray irradiation: (**g**) γH2AX; (**h**) 53BP1. Scale bar, 25 μm

## Discussion

HGPS modeling has traditionally relied on patient-derived fibroblasts, which have revealed key cellular hallmarks such as nuclear envelope abnormalities ^50^, persistent DNA damage ^51,52^, and epigenetic defects ^53,54^. However, these models do not recapitulate the tissue-specific pathology of HGPS, which primarily affects mesoderm-derived lineages (e.g., vascular smooth muscle, skeletal muscle, and bone) while sparing ectodermal tissues like neurons. To overcome these limitations, iPSCs reprogrammed from HGPS patient fibroblasts have been utilized to derive a number of cell types of interest, including vascular smooth muscle cells, mesenchymal stem cells, and neural cells ^21,22,55^.

Herein, we established two isogenic pairs of hPSCs differing only at the *LMNA* locus and differentiated them into NMOs—a 3D multicellular platform that recapitulates both mesoderm-derived muscle and ectoderm-derived neural lineages (Fig. 1) and thereby uniquely enables side-by-side evaluation to clearly reveal mesoderm-specific pathologies and transcriptome signatures under a shared genetic background (Fig. 2). Strikingly, we found that perinuclear progerin accumulation and associated pathologies, including sequestration of 53BP1 and defective recruitment of key proteins such as γH2AX, 53BP1, and ATM at DNA damage repair foci, are exclusive to the muscle compartment (Fig. 3). This mirrors the selective vulnerability observed in vivo in mesoderm-derived tissues such as vasculature, bone, and muscle, a vulnerability not echoed in neural lineages. Interestingly, the selective absence of progerin accumulation in the neural compartment of our NMOs, which recapitulates the resistant HGPS pathology, offers a unique opportunity to investigate the endogenous mechanisms that restrict progerin production in neural tissues as seen in iPSC-derived neural cells ^55^.

Using our NMO model, wherein perinuclear progerin accumulation, loss of DNA damage foci, and even loss of perinuclear heterochromatin structure all serves as robust pathogenic readouts, we successfully demonstrated the therapeutic efficacy of FATE, a precision base editing strategy designed to disrupt the CSIM farnesylation motif of LMNA (Fig. 5). Unlike previous base editing approaches that target the canonical c.1824C>T (p.G608G) mutation found in classical HGPS ^16^, this mutation-agnostic editing strategy can apply to a broader spectrum of *LMNA* mutations, including the non-classical *LMNA* variants c.1864C>G (p.T623S), c.1940T>G (p.L647R), and c.1968+1G>A, as well as ZMPSTE24 loss-of-function laminopathies, all of which similarly result in aberrantly farnesylated, uncleaved prelamin A. Crucially, the only FDA-approved drug for HGPS patients, lonafarnib, broadly blocks farnesylation across numerous essential proteins, including small GTPases and mitotic regulators. Distinct from that nonselective approach, FATE introduces a single-base substitution specifically within the codon of *LMNA* that encodes the cysteine of the CSIM motif. Whole-exome sequencing confirmed the editing to be confined to the intended *LMNA* locus, with no off-target changes detected in other genes encoding farnesylated proteins (e.g., *LMNB1*, *LMNB2*, *HRAS*, *NRAS*, *KRAS4A/B*, *RHOE*, *RHEB*, *CENPE*, or *CENPF*) (Fig. 4). This exceptional specificity highlights the potential of FATE as a targeted and safe genome editing strategy by which to eliminate pathogenic farnesylation without perturbing the global prenylation landscape. Notably, prior studies have reported conflicting findings on the toxicity of non-farnesylated progerin (SSIM vs. CSM) ^56,57^, the basis of which remains unresolved.

Prior studies in HGPS mouse models harboring the human-equivalent L*mna* mutation (c.1827C>T; p.G609G) ^19^, as well as models carrying a human LMNA gene fragment encompassing the pathogenic c.1824C>T site ^16,58^, have established foundational strategies for in vivo therapeutic gene editing. Distinct from these typically mutation-specific approaches, however, FATE targets a shared pathogenic post-translational modification—farnesylation—and therefore represents a broadly applicable, mutation-agnostic therapeutic platform for diverse HGPS variants. Building on this concept, we demonstrated that mRNA-based delivery of FATE rescues disease phenotypes in neuromuscular organoids derived from *LMNA*-mutant hPSCs, establishing proof-of-concept for the *in situ* correction of HGPS pathology. Notably, unlike adeno-associated-virus–based approaches, FATE mRNA delivery is compatible with repeat dosing—a critical advantage for addressing the progressive and cumulative nature of HGPS. Collectively, the results of this work provide a strong rationale for future studies employing LNP–mediated FATE delivery in HGPS animal models to assess tissue-specific editing efficiency, durability of correction, and long-term safety.

## Methods

### Culture of human pluripotent stem cells

Two human pluripotent stem cell lines were used in this study: HGPS patient-derived iPSCs (the following cell line was obtained from the NIGMS Human Genetic Cell Repository at the Coriell Institute for Medical Research: AG28340; referred to as HGPS-iPSCs) and the human embryonic stem cell line WA09 (H9; obtained from WiCell Research Institute, Madison, WI, USA; referred to as WT-hESCs). To generate isogenic controls, the c.1824C>T mutation in HGPS-iPSCs was corrected using the PE5stem prime editing system as described by Park et al. (Nat Commun, 2024), resulting in mutation-corrected Edit-iPSCs. Conversely, the same c.1824C>T mutation was introduced into WT-hESCs using the same system to generate Mutant-iPSCs. Two sets of human pluripotent stem cells (hPSCs) were maintained in mTeSR™1 medium (STEMCELL Technologies) on Matrigel-coated plates (Corning, 354277) at 37 °C in a humidified incubator with 5% CO . All hPSC lines used in this study were tested and confirmed negative for mycoplasma contamination. Cells were passaged approximately every four days using Versene solution (Thermo Fisher Scientific).

### FATE plasmid construction

To generate FATE-iPSCs, the px330-mCherry plasmid (Addgene #98750, a gift from Dr. Jinsong Li) was modified by replacing the Cas9 coding sequence with the SpRY-ABE8e variant from pCMV-SpRY-ABE8e (Addgene #185671, a gift from Dr. David Liu) using Gibson assembly. Following this, a guide RNA sequence (5’-TGCTGCAGTTCTGGGGGCTC-3’) was inserted into the BbsI restriction site of the modified vector.

### Generation of neuromuscular organoids (NMOs) from hPSCs via induction of human neuromesodermal progenitors (hNMPs)

Human pluripotent stem cells (hPSCs) were dissociated into single cells using Accutase (STEMCELL Technologies) when cultures reached approximately 70–80% confluency. Cell counts were performed using the Countess III automated cell counter (Thermo Fisher), with the cell size threshold set to include only particles with a diameter ≥9 μm. Based on growth characteristics of each cell line, 1.3–1.5 × 10 cells were seeded per well for WT-hESCs and Mutant-iPSCs, whereas 1.0–1.2 × 10 cells were used for HGPS-iPSCs and Edit-iPSCs. All cells were plated onto Matrigel-coated (Corning, 354277) 6-well plates. On day 0, to reduce stress from cell dissociation, 10 μM Y-27632 (a ROCK inhibitor) was added to the NMO basal medium, which also included 3 μM CHIR99021 and 40 ng/ml basic fibroblast growth factor (bFGF). The NMO basal medium was composed of a 1:1 blend of Advanced DMEM/F12 (supplemented with 1× N2) and Neurobasal medium (supplemented with 1× B27, 2 mM L-glutamine, 75 μg/ml BSA Fraction V, and 0.1 mM 2-mercaptoethanol). On day 1, fresh medium of the same composition—except without Y-27632—was applied, and daily medium changes were performed until day 3. This 2D induction protocol facilitated the differentiation of hPSCs into neuromesodermal progenitors (NMPs).

On day 3, NMPs were enzymatically dissociated into single cells using Accutase and seeded at a density of 8,000 cells per well into ultra-low attachment 96-well plates. Cell counts were obtained using a Countess III automated cell counter, and aggregation was initiated by centrifugation at 350 × g for 2 minutes. For organoid formation (day 0 of 3D culture), 100 μl of NMO basal medium supplemented with 50 μM Y-27632, 10 ng/ml bFGF, 2 ng/ml IGF1, and 2 ng/ml HGF was added to each well. On day 2, 50 μl of medium was carefully removed and replaced with 100 μl of fresh NMO basal medium containing 2 ng/ml IGF1 and 2 ng/ml HGF. On day 4, the medium was completely replaced with NMO basal medium without additional growth factors and subsequently refreshed every other day. On day 10, organoids were transferred into 60-mm ultra-low attachment dishes containing 5 ml of NMO basal medium, and on day 30, they were moved to 100-mm Petri dishes with 12 ml of the same medium. Medium changes continued every two days. To minimize mechanical stress, pipette tips were trimmed using autoclaved scissors during transfers. From day 10 onward, organoids were maintained on an orbital shaker set to 75 rpm to facilitate maturation.

### Immunohistochemistry of organoids

Organoids were collected at specific time points, rinsed once in DPBS, and fixed in 4% paraformaldehyde (PFA) at 4°C for 1 to 2 hours depending on the duration of culture. Post-fixation, the samples were washed three times in DPBS and placed in 30% sucrose at 4°C overnight, allowing them to sink completely as an indication of full infiltration. For embedding, organoids were transferred to molds filled with Tissue-Tek O.C.T. compound. Cryopreservation was achieved by partially solidifying isopentane (2-methylbutane) in a metal container surrounded by liquid nitrogen in a styrofoam box. Once the isopentane appeared opaque, the molds were immediately submerged to freeze the samples. The frozen blocks were stored at –80°C prior to cryosectioning. Tissue sections of 10 μm were sliced using a Leica CM1520 cryostat maintained at –19°C and mounted onto glass slides. Before immunostaining, slides were equilibrated to room temperature and treated with 0.3% Triton X-100 in DPBS to eliminate any remaining O.C.T. compound. Non-specific binding was blocked for 1 hour using either 5% normal goat serum or 4% BSA in DPBS. Sections were then incubated overnight (12-16h) at 4°C with primary antibodies diluted in the blocking buffer. The following day, sections were washed three times and treated with fluorescently labeled secondary antibodies for 1 to 1.5 hours at room temperature. Nuclei were stained with DAPI (1 μg/ml) for 10 minutes and rinsed again. Final mounting was done using ProLong™ Gold Antifade Reagent. All 2D images were acquired using a Leica SP8 confocal microscope. For 3D confocal imaging, samples were imaged with an LSM980 confocal microscope, and rotational movies (360° along the Y-axis at 250 fps) were generated using the Zen Blue software.

### Real-Time PCR analysis

Easy-BLUETM RNA isolation kit (iNtRON Biotechnology) was used for total RNA extraction. For reverse transcription, 5× PrimeScriptTM RT mix (TaKaRa) was used to generate cDNA. Quantitative real-time PCR was performed with SYBR Green PCR reagents (Life Technologies) using the QuantStudio3 (Applied Biosystems).

### Immunoblotting analysis

Organoids were homogenized in RIPA buffer (Biosesang) supplemented with 1 μM protease inhibitor and 10 μM sodium orthovanadate using a TissueLyser II (QIAGEN) set to a frequency of 30 Hz for 3 minutes. The homogenates were extracted on ice for 2 hours with intermittent vortexing every 10 minutes. Lysates were then centrifuged at 13,000 rpm for 20 minutes at 4 °C, and the resulting supernatants were collected as total protein lysates. Protein concentrations were determined using the Pierce™ BCA Protein Assay Kit (#23225, Thermo Fisher Scientific). Equal amounts of protein were diluted with Bolt™ LDS Sample Buffer (4X) and Bolt™ Sample Reducing Agent (10X) to achieve uniform concentrations and then boiled at 95 °C for 10 minutes.

Following protein extraction, immunoblotting was carried out using the following protocol. Approximately 20–30 μg of protein per sample was loaded onto 3–8% NuPAGE™ Tris-Acetate Mini Protein Gels (Invitrogen) and separated by electrophoresis at 150 V. The resolved proteins were transferred onto PVDF membranes using iBlot™ 3 Transfer Stacks (MIDI, PVDF format) via dry transfer. Membranes were blocked in 5% bovine serum albumin (BSA) in TBS-T for 1–2 hours at room temperature and then incubated overnight at 4 °C with primary antibodies diluted in the same blocking solution. After three washes with TBS-T, membranes were incubated with HRP-conjugated secondary antibodies (1:10,000 dilution in TBS-T) for 1–1.5 hours at room temperature. Signal detection was performed using either Miracle-Star (#16028, iNtRON Biotechnology) or Amersham™ ECL Prime (#GERPN2232, Cytiva) chemiluminescence reagents.

### Fractionation of organoids and immunoblotting analysis

Nuclear membrane fractionation was performed based on the protocol described by Udi et al. (J Cell Biol, 2023). Briefly, 4-6 organoids were dissociated into single cells using Liberase™ TL Research Grade (Roche) following the manufacturer’s instructions. The resulting cell suspension was passed through a Falcon® 40 μm strainer to remove aggregates and debris. Cells were pelleted by centrifugation at 300 × g for 3 minutes, washed once with 1 ml pre- chilled PBS, and pelleted again under the same conditions. After gently removing the supernatant, the cell pellet was resuspended in lysis buffer (at a 1:5 to 1:8 v/v ratio), followed by pipetting and brief vortexing. The suspension was incubated on ice for 5 minutes to allow swelling. Cells were then lysed mechanically using a 26G 1/2” KOVAX-SYRINGE. The lysate was carefully underlaid with 200 μl of 20% sucrose in 8% PVP buffer and centrifuged at 3,500–4,000 × g for 10 minutes at 4 °C. The supernatant, including the sucrose phase, was collected as the cytoplasmic fraction. The pelleted nuclei were resuspended in DNase buffer (1 ml per 1×10 cells), vortexed, and incubated at room temperature for 30 minutes. The suspension was then underlaid with 100 μl of 30% sucrose prepared in 20 mM HEPES (pH 8), 0.1 mM MgCl, and 1:100 Solution P. Nuclear envelopes were pelleted by centrifugation at 18,000 × g for 10 minutes at 4 °C. The resulting supernatant was collected as the nucleoplasmic fraction, while the pellet represented the nuclear membrane fraction.

For downstream SDS-PAGE and Western blot analysis, each fraction was processed by methanol precipitation. Specifically, 900 μl of methanol was added to 100 μl of sample, followed by vortexing and overnight incubation at –20 °C. Samples were centrifuged at 20,000 × g for 20 minutes at 4 °C, and pellets were briefly sonicated (2 seconds) in 500 μl of methanol, incubated again at –20 °C for 20 minutes, and re-pelleted. The final pellet was resuspended in a 1× dilution of Bolt™ LDS Sample Buffer (4X) and Bolt™ Sample Reducing Agent (10X) for protein denaturation.

#### 8% PVP Buffer

This buffer was composed of 8% polyvinylpyrrolidone (PVP-40), 20 mM potassium phosphate, and 7.5 μM magnesium chloride (MgCl). The pH was adjusted to 6.53 using concentrated phosphoric acid (H PO).

#### Lysis Buffer

The lysis solution contained 6% PVP-40, 0.015% Digitonin, 0.015% Triton X-100, and 4 μM Cytochalasin B. Protease inhibitors were added at a final dilution of 1:100 from Solution P. Dithiothreitol (DTT) was included at 1 mM final concentration

#### DNase Buffer

DNase treatment was performed using a buffer consisting of 10% sucrose in 20 mM HEPES (pH 8.0), supplemented with 0.1 mM MgCl, 10 μM CaCl, 1 mM DTT, 100 μg/mL heparin, and 0.1 μg/mL RNase A. DNase I (Roche) was added at 1:1,000 dilution from a 5 mg/mL stock prepared in 50% glycerol containing 10 mM Tris-HCl (pH 7.4), 50 mM NaCl, 1 mM DTT, and 2 mM MgCl . Protease inhibitors were also added at 1:100 dilution from Solution P.

### Electron microscopy of organoids

Organoids were fixed overnight at 4 °C in Karnovsky’s fixative, consisting of 2% (w/v) paraformaldehyde and 2% (v/v) glutaraldehyde in 0.05 M sodium cacodylate buffer(pH 7.4). Following fixation, samples were washed three times for 5 minutes each at room temperature in 0.05 M sodium cacodylate buffer. Post-fixation was performed using 1% osmium tetroxide diluted in 0.1 M sodium cacodylate buffer. The samples were then rinsed three times with distilled water at room temperature, each for 5 minutes, and stained en bloc overnight at 4 °C in 0.5% uranyl acetate diluted in distilled water.

After En Bloc staining, samples were again washed three times with distilled water for 5 minutes each and dehydrated through a graded ethanol series (30%, 50%, 70%, 80%, 90%, and 100%). Dehydrated samples were treated twice with 100% propylene oxide at room temperature for 15 minutes per wash. Resin infiltration was performed sequentially using mixtures of propylene oxide and Spurr’s resin at 1:1 and 1:2 ratios, each for 1.5 hours at room temperature, followed by transition into 100% Spurr’s resin overnight. Finally, samples were embedded in fresh 100% Spurr’s resin in embedding molds and polymerized at 70 °C overnight.

Ultrathin sections (∼70 nm) were stained with 2% uranyl acetate followed by 3% lead citrate. Imaging was performed using a JEM-1010 transmission electron microscope (JEOL Ltd., Tokyo, Japan) operated at 80 kV. Images were acquired using a CCD camera equipped with an image capture system and processed with Radius 2.1 software (EMSIS GmbH, Münster, Germany).

### Single cell preparation from organoids for scRNAseq

For single-cell RNA sequencing (scRNA-seq), 6 of day 70 organoids (HGPS-NMOs, Edit-NMOs, and FATE-NMOs) were washed with DPBS and transferred to 1.5 mL microcentrifuge tubes. Organoids were then incubated with 1 mL of Liberase™ TL Research Grade (Roche) at 37°C for 30 minutes, following the manufacturer’s protocol. After enzymatic digestion, mechanical dissociation was performed by pipetting until a single-cell suspension was achieved. The suspension was passed through a 40 μm cell strainer (Falcon®) to eliminate cell aggregates and debris. The resulting cell suspension was washed with DPBS and resuspended in HypoThermosol® FRS (STEMCELL Technologies), followed by quality assessment using the Celinius single-cell QC platform.

### Sample Preparation and Quality Control

Single-cell RNA sequencing (scRNA-seq) was performed using the 10x Genomics Chromium platform. High-quality Dissociated neuromuscular organoid cells were prepared to ensure optimal assay performance. All suspensions were washed and resuspended in 1 x PBS containing 0.04% BSA buffer, using wide-bore pipette tips to prevent shearing. The quality of the final suspension was assessed by evaluating cell concentration, viability, and the degree of aggregation to ensure suitability for microfluidic partitioning.

### scRNA-seq Library Preparation and Sequencing

Single cells were partitioned into Gel Beads-in-emulsion (GEMs) using a Chromium controller Within each GEM, mRNA was captured and barcoded. scRNA-seq libraries were subsequently generated following the manufacturer’s protocol (10x genomics, CG000315), which includes reverse transcription, cDNA amplification, and library construction Chromium Next GEM Single Cell 3’ Kit v3.1 (10x Genomics, PN1000268) The quality and size distribution of the resulting cDNA and final libraries were assessed, and concentrations were quantified. Libraries were sequenced on a compatible platform using NovaSeq 6000 platform(Illumina) and a paired-end configuration, with read lengths structured to capture the cell barcode, Unique Molecular Identifier (UMI), and the transcript sequence.

### Primary Data Processing

Raw sequencing base call (BCL) files were demultiplexed into FASTQ files using the cellranger mkfastq pipeline. The resulting FASTQ files were processed with cellranger (10x Genomics, v-8.0.1)count to align reads to a GRCm38 reference genome and generate a feature-barcode count matrix. This matrix, containing UMI counts per gene for each cell, served as the input for all subsequent analyses. All processing was performed using a consistent version of the Cell Ranger software to ensure reproducibility.

### Single-cell RNA-seq data analysis

Single-cell RNA-seq datasets from HGPS, EDIT, and FATE samples were processed using Seurat v4.3.0 (Stuart et al., 2019). Cells with fewer than 200 detected genes or with more than 15% mitochondrial transcripts were removed. Normalization and variance stabilization were performed using an SCTransform-based approach, and the datasets were integrated after selecting 3000 highly variable features. Batch effects were corrected during the integration step. The integrated data were dimensionally reduced with principal component analysis (PCA), and the first 15 principal components were used for constructing a shared nearest neighbor graph and subsequent clustering. Low-dimensional visualization was performed with UMAP to explore cellular heterogeneity. Cell clusters were annotated into five major lineages (skeletal muscle, neuron, sclerotome, endothelial, and epithelial) based on canonical marker gene expression and module score analysis of curated gene sets . For cross-sample comparison, reference-based label transfer was applied to map the FATE dataset onto the integrated HGPS reference, and the query cells were projected onto the existing UMAP embedding.

### Differential expression and pathway enrichment analysis

For lineage-specific comparisons, skeletal muscle and neuronal clusters were separately subsetted from the integrated HGPS and EDIT dataset to generate SM (skeletal muscle) and NR (neuron) objects. SCTransform normalization was applied independently to each subset. Differentially expressed genes (DEGs) between HGPS and EDIT were identified using a log2 fold change threshold > 1 and adjusted p-value < 0.05. Over-representation analysis of DEGs was conducted using EnrichR (Kuleshov et al., 2016) with the WikiPathways_2024_Human database to identify lineage-specific functional pathways. Additionally, gene set enrichment analysis (GSEA) was performed using the fgsea package (Korotkevich et al., 2021) with cell type signatures from the MSigDB C8 collection (via msigdbr). Ranked gene lists based on log2 fold change were tested for enrichment, and significant pathways were defined by normalized enrichment scores (NES) with adjusted p-values < 0.05. Directionality of enrichment was interpreted based on the sign of NES, with positive and negative values indicating enrichment in HGPS and EDIT, respectively. Enrichment results were visualized by plotting NES against –log10 adjusted p-values.

### Pathway Activity Estimation and PCA Analysis Based on Pseudobulk GSVA

To assess pathway activity at the group level, pseudobulk expression matrices were generated by averaging SCTransform-normalized expression values per condition and cell type (skeletal muscle and neuron) from both the integrated (HGPS and EDIT) and mapped (FATE) datasets. Gene Set Variation Analysis (GSVA) was conducted using the GSVA R package (Hänzelmann et al., 2013) with curated gene sets from the KEGG Legacy and Human Phenotype Ontology collections, with emphasis on DNA damage repair pathways. GSVA scores representing relative pathway activity were computed for each pseudobulk sample. These scores were used for downstream comparative analysis, including hierarchical clustering and principal component analysis (PCA). PCA was performed on the GSVA enrichment matrix derived from KEGG Legacy pathways to evaluate global pathway activity patterns across HGPS, EDIT, and FATE samples.

### Derivation and Application of Young Muscle Signature

To derive a transcriptional signature representing youthful skeletal muscle, we utilized publicly available human muscle single-cell RNA-seq data from Veronika R. K. et al. (Nature Aging, 2024). Donor samples were stratified into young (ages 15–40) and old (ages 50–60 and 50–75) cohorts. Differential expression analysis was performed between the two groups, and genes with adjusted p-value < 0.05 and average log2 fold change > 1 were selected as significantly upregulated in the young group. These genes were defined as the young muscle signature.

To evaluate this signature in experimental samples, pseudobulk profiles from myogenic progenitor cells in the HGPS, EDIT, and FATE datasets were generated. GSVA was then applied to quantify the enrichment of the young muscle signature, enabling assessment of youthful transcriptional programs across conditions.

### Whole Exome Sequencing (WES) and Data Processing

Whole exome capture libraries were prepared using the Agilent SureSelect Target Enrichment protocol for Illumina paired-end sequencing libraries (Version C2, December 2018) with 0.5 μg of input genomic DNA. The SureSelect Human All Exon V8+UTR probe set was used in all cases. DNA quality and quantity were assessed with PicoGreen (Invitrogen) and Agilent TapeStation gDNA screentape. One microgram of genomic DNA was diluted in EB buffer and sheared to a target peak size of 150–200 bp using a Covaris LE220 focused ultrasonicator (Covaris, Woburn, MA) according to the manufacturer’s recommendations. Fragmented DNA underwent end repair, A-tailing, and adapter ligation, followed by PCR amplification. For exome capture, 250 ng of DNA library was hybridized with the capture probes at 65 °C for 24 h using the heated lid option (105 °C) on a thermal cycler, washed, and PCR amplified. Final libraries were quantified using qPCR (KAPA Library Quantification Kit for Illumina platforms) and qualified on an Agilent TapeStation D1000 screentape. Sequencing was performed on the Illumina NovaSeqX platform (Illumina, San Diego, USA) with a target coverage depth of approximately 400×, yielding an average achieved coverage of ∼250×. Raw sequencing reads in FASTQ format were subjected to initial quality control to assess read quality, adapter content, and duplication rates. Reads were aligned to the human reference genome (GRCh38) using BWA-MEM, and PCR duplicates were removed with Picard MarkDuplicates. Base quality score recalibration (BQSR) was performed using GATK BaseRecalibrator, and the resulting BAM files were processed following GATK Best Practices. Variant calling was carried out using GATK HaplotypeCaller in GVCF mode, followed by joint genotyping with GenotypeGVCFs. Variants were filtered based on quality metrics with GATK VariantFiltration and functionally annotated using SnpEff and dbNSFP. Final results were compiled into both VCF and tabular formats for downstream analyses.

### Targeted Deep Sequencing and Data Processing

Genomic DNA from induced pluripotent stem cells (iPSCs) and nuclear membrane organoids (NMOs) was extracted using the Wizard Genomic DNA Purification Kit (#A1120, Promega) to evaluate target editing efficiency. The target locus was amplified using Solg™ Pfu-X DNA Polymerase (#SPX16-R25h, Solgent) to generate sequencing libraries. Libraries were sequenced on an Illumina MiSeq platform (Binonics, South Korea) with 150-bp paired-end reads at a depth of ∼30,000 reads. FASTQ files were aligned to the human reference genome (hg38) using BWA-MEM2, and the resulting SAM files were converted to BAM format, sorted, and indexed using SAMtools for downstream analysis.

### Identification of gRNA-dependent Off-target Effects

To identify potential off-target effects associated with the designed guide RNA (gRNA), we first predicted candidate loci with high sequence similarity (0–2 mismatches) using RgenTools. A total of 37 candidate sites were identified. Among these, 19 were located within protein-coding regions, including 5 within exons, while the remainder were distributed across intronic and intergenic regions. To assess editing events at these loci, we applied region-specific data sources:

- For exonic sites, we analyzed whole-exome sequencing (WES) data.
- For intronic and intergenic sites, we used Targeted deep sequencing data.

In both cases, aligned BAM files were processed to extract base-level read information, and conversion rates were calculated by dividing the number of reads supporting the expected substitution by the total read depth at each site. Only substitutions with frequencies exceeding the estimated sequencing error rate (∼1%) were considered indicative of potential off-target editing.

### Identification of gRNA-independent Off-target Effects

To identify sequence alterations not attributable to gRNA targeting, we analyzed VCF files derived from WES data. Positions with a read depth >200 were retained for further inspection. Sites showing conversion in both Mock and FATE samples were excluded to focus on treatment-specific effects. As a result, six candidate sites, including one in LMNA, were uniquely observed in FATE-treated samples, while nine sites were identified only in Mock-treated samples.

### Analysis of CAAX Motif Sequences

To examine the conservation of the CAAX motif, we extracted 12-nucleotide sequences corresponding to CAAX regions from genes known to harbor this motif, including LMNA, small GTPases, lamins, and the CENP gene family. Genomic coordinates of CAAX motifs were used to retrieve base-level read data from WES BAM files. Nucleotide frequencies at each position were aggregated across all selected motifs, and a sequence logo was generated to visualize base preferences and positional conservation within the motif.

### mRNA synthesis via *In vitro* transcription (IVT)

Linear DNA templates were generated from the plasmid pCMV-SpRY-ABE8e (Addgene #185671) by PCR with PrimeSTAR® HS DNA Polymerase (Takara) following the manufacturer’s protocol. A 120-nt poly(A) tail was encoded in the template using a poly(T)-containing primer during PCR. The PCR products were purified using AccuPrep® PCR/Gel Purification Kit (Bioneer) and verified by agarose gel electrophoresis (1% agarose in 1× TBE buffer).

*In vitro* transcription was performed using MEGAscript™ T7 Transcription Kit (Thermo Fisher Scientific), with CleanCap® AG (TriLink) and substituting UTP with N1-methyl-pseudouridine-5′-triphosphate (N1-Me-ΨUTP). Transcription reactions were carried out at 37 °C for 2 h, followed by DNase treatment to digest any residual template DNA. The resulting mRNA was purified using HiGene™ DNA/RNA Purification Mini Column (Biofact). Transcript integrity was confirmed by denaturing agarose gel electrophoresis (1% agarose in 1× MOPS buffer).

### Preparation and Characterization of LNPs

LNPs were prepared using a microfluidic mixing system (Ignite, Precision NanoSystems). A lipid mixture containing an ionizable lipid (undisclosed structure), 1,2-distearoyl-sn-glycero-3-phosphocholine (DSPC), cholesterol, and PEG-lipid were dissolved in ethanol at the specified molar ratio. The ethanol phase (97.5% v/v) was mixed with 10 mM citrate buffer, pH 3.0 (2.5% v/v). ABE mRNA and sgRNA were diluted in an aqueous phase (1× PBS and 10 mM citrate buffer, v/v = 2:1). LNPs were formulated by microfluidic mixing at a total flow rate of 12 mL/min with an ethanol-to-aqueous phase ratio of 1:3. The resulting LNPs were diluted 40-fold in 1× PBS and concentrated using Amicon® Ultra-15 centrifugal filters, 10 kDa MWCO (Millipore).

Particle size and polydispersity index (PDI) of LNPs were measured by dynamic light scattering (DLS) using a Zetasizer Lab Blue (Malvern Panalytical). RNA encapsulation efficiency was quantified with Quant-iT™ RiboGreen® RNA Assay Kit (Thermo Fisher Scientific) as described below.

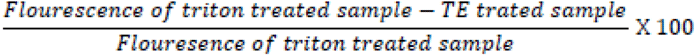

### mRNA-LNP microinjection

For FATE mRNA–LNP delivery into NMOs, organoids were transferred into a 100 μL basal medium drop using a micropipette and moved on the micromanipulator stage. A microinjector (CellTram, Eppendorf) connected to a non-heparinized microhematocrit capillary tube (DWK Life Sciences) was used as a holding pipette to hold the neural region of the organoid in place. A second microinjector (FemtoJet 4i, Eppendorf) was then used to perform three injections at a 90° angle into the muscle region under the following conditions: injection pressure of 150-250 hPa (adjusted by confirmation of LNP entry into the organoid), and consistent pressure of 0 hPa. To prevent backflow of the injected LNP solution from organoids, the injection needle was kept inside the organoid for 3 seconds before withdrawal. Following injection, organoids were transferred to NMO basal medium using a 1-ml pipette tip with the end trimmed and subsequently cultured for 10 days in 60-mm ultra-low attachment plates prior to NGS analysis.

### Calculation of Editing Efficiency via mRNA Delivery

To evaluate genome editing outcomes following mRNA delivery, Targeted deep sequencing data were analyzed as described above. Base composition at the editing site was quantified from the aligned reads, and the editing frequency was calculated using the same threshold-based approach. This allowed direct estimation of editing efficiency at the nucleotide level.

### Data and Code Availability

Whole exome sequencing (WES) data have been deposited in the Sequence Read Archive (SRA) under the accession number PRJNA1306877. Three single-cell RNA sequencing datasets are available in the Gene Expression Omnibus (GEO) under the accession number GSE305237. The R and Shell scripts used for the analysis are available from the Lead Contact upon request.

## Acknowledgements

This research was supported by a grant of the Korea-US Collaborative Research Fund (KUCRF) (RS-2024-00466703), ABC-based Regenerative BioTherapeutics (ABC project) grant (RS-2024-00432867) and National Research Foundation of Korea (NRF) grant (RS-2025-00560943), funded by the Ministry of Science and ICT and Ministry of Health & Welfare, Republic of Korea.

## Author information

These authors contributed equally: Dong-Woo Kim, Eun-Ji Kwon

Authors and Affiliations

**College of Pharmacy, Seoul National University, Seoul, Republic of Korea**

Dong-Woo Kim, Eun-Ji Kwon, Hyukjin Lee & Hyuk-Jin Cha

**Research Institute of Pharmaceutical Sciences, Seoul National University, Seoul, Republic of Korea**

Hyukjin Lee & Hyuk-Jin Cha

**Laboratory of Theriogenology and Biotechnology, Department of Veterinary Clinical Science, College of Veterinary Medicine and the Research Institute of Veterinary Science, Seoul National University, Seoul, Republic of Korea**

Beom-Jin Jeon, Dong-Hyeok Kwon & Goo Jang

**College of Pharmacy, Graduate School of Pharmaceutical Sciences, Ewha Womans University, Seoul 03760, Republic of Korea**

Youngyoon Yoon & Yuna Hwang

## Contributions

HJ.C conceived the overall study design, led the experiments, and wrote the manuscript. DW.K conducted experiments, engaged in critical discussions of the results, and wrote the initial draft of the manuscript. EJ.K performed scRNAseq analysis and in silico analysis and wrote the initial draft of the manuscript. BJ.J DH.K and G.J conducted the gene delivery and sample preparation. H.L provided mRNA and LNP. All authors contributed to writing and revising the manuscript and approved the final version.

## Corresponding authors

Correspondence to Hyuk-Jin Cha

## Ethics declarations

Competing interest

The authors declare that they have no competing interests.

