## Supplementary Figures and Video for "Mutation-Agnostic Base Editing of the Progerin Farnesylation Site Rescues Hutchinson-Gilford Progeria Syndrome Phenotypes in Neuromuscular Organoids": Inventory of Supporting Information.pdf

**Running Title:** Mutation-Agnostic Editing of Lamin A Farnesylation in Progeria

Dong-Woo Kim*, Eun-Ji Kwon*, Beom-Jin Jeon, Dong-Hyeok Kwon, Goo Jang, Youngyoon Yoon, Yuna Hwang, Hyukjin Lee and Hyuk-Jin Cha#

#### Extended Data Figure Legends

**Extended Data Fig. 1: Lamin A expression in two sets of hPSCs and establishment of two sets of isogenic NMO pairs.**

**a**, Real-time PCR analysis of LMNA (Lamin A), GAPDH as an internal control. **b**, Immunofluorescence images showing established NMOs derived from two sets of isogenic hPSCs at day 40 and day 70. At day 40, fast myosin heavy chain (magenta, muscle marker) and Tuj1 (green, neuron marker) are shown. At day 70, Ttn (green, muscle marker) and GFAP (red, glial marker) are shown. Scale bars, 200 μm (day 40) and 500 μm (day 70). **c**, Transmission electron microscopy images of day 70 HGPS-, Edit-, Mutant-, and WT-NMOs showing two major functional compartments. The muscle region displays advanced sarcomeric organization, including aligned myofibrils with distinct Z-lines (Z) and M-bands (M). The neural region reveals neurofilaments (NF), mitochondria (Mi), and synaptic vesicles (SV), indicative of mature synaptic structures. Scale bars are indicated in each panel. **d**, Immunofluorescence images of day 40 Mutant- and WT-NMOs stained with Progerin (green), Desmin (magenta), and DAPI (blue). Scale bar, 25 μm.

**Extended Data Fig. 2: Single-cell analysis of neural cellular composition and developmental trajectories in HGPS and Edit NMOs.**

**a**, Sub-clustering of the neural trajectories reveals 5 distinct subpopulations. Shades of orange and pink indicate neural subclusters. Relative proportions of each subcluster in HGPS organoids are shown below. **b**, Dot plot showing representative gene expression across three neural subclusters. **c**, Human Phenotype Ontology terms enriched in HGPS neural cells compared to Edit controls. **d**, Pathway enrichment analysis of neuronal compartments in HGPS- and Edit-NMOs.

**Extended Data Fig. 3: Single-cell analysis and altered 53BP1 localization in HGPS-NMOs**

**a**, Heatmap showing GSVA-based KEGG pathway enrichment in neuronal and muscle compartments of HGPS- and Edit-NMOs. Gene sets related to metabolism are highlighted in red, and those related to autophagy are highlighted in blue. **b**, Schematic illustrating the heterozygous expression of LMNA and Progerin in HGPS iPSCs, driven by a shared promoter resulting in a 1:1 expression ratio. **c**, Cells with SRSF1 and SRSF2 expression in the top 25% were designated as the "SRSF1/ SRSF2-high" group (top). UMAP visualization shows the distribution of SRSF1/ SRSF2-high and SRSF1/ SRSF2-low groups across organoids (middle). Heatmap displaying enrichment of DNA repair–related gene sets in SRSF1/ SRSF2-high and SRSF1/ SRSF2-low groups within the neuronal and muscle compartments of HGPS and Edit organoids (bottom). **d**, Immunofluorescence images of day 50 HGPS and WT NMOs stained with 53BP1 (green), and DAPI (blue), showing both neuronal and muscle regions. Scale bar, 25 μm.

**Extended Data Fig. 4: Impaired foci formation of DNA damage repair machinery specifically in the muscular compartment of HGPS and mutant NMOs.**

**a**, Immunofluorescence images of day 70 HGPS and Edit NMOs stained with Progerin (green), γH2AX (red), and DAPI (blue), showing the neuronal region under mock, 1-hour, and 4-hour conditions following 5 Gy X-ray irradiation. Scale bar, 25 μm. **b**, Immunofluorescence images of day 70 HGPS and Edit NMOs stained with Progerin (green), pATM (red), and DAPI (blue), showing the muscle region under mock and 1-hour conditions following 5 Gy X-ray irradiation. Scale bar, 25 μm. **c, d**, Immunofluorescence images of day 70 Mutant and WT NMOs stained with Progerin (green), γH2AX (red), and DAPI (blue), showing both muscle and neuronal regions under mock and 1-hour conditions following 5 Gy X-ray irradiation. Scale bar, 25 μm. **e**, Immunofluorescence images of day 70 mutant and WT NMOs stained with Progerin (green), pATM (S1981) (red), and DAPI (blue), showing the muscle region under mock and 1-hour conditions following 5 Gy X-ray irradiation. Scale bar, 25 μm.

**Extended Data Fig. 5: Heterochromatin organization in the neural compartment of isogenic NMOs derived from HGPS-iPSCs and WT-hESCs.**

**a**, Transmission electron microscopy (TEM) images showing the nuclear heterochromatin structure in the neural region of neuromuscular organoids (NMOs) derived from two isogenic HGPS hPSC pairs.

**Extended Data Fig. 6: Atypical HGPS variants, technical challenges, and FATE strategy with gRNA design.**

**a**, Table summarizing classical and non-classical HGPS mutation types, including the mutation position, example protospacer sequences for sgRNA design, the corresponding genome editing tools (ABE or PE or CGBE), and associated concerns such as bystander editing or transversion mutations. **b**, Immunofluorescence images of cells overexpressing Progerin-GFP, Progerin (SSIM)-GFP with a C661S mutation. **c**, Immunofluorescence images of cells overexpressing Progerin-GFP or Progerin (SSIM)-GFP with a C661S mutation after 5 Gy X-ray irradiation, stained with 53BP1 (red) and DAPI (blue). Scale bar, 25 μm. **d**, Table showing candidate sgRNA sequences, PAMs, and compatible ABE versions for targeting the CSIM motif in exon 12 of the LMNA gene for FATE. The schematic above depicts the partial LMNA locus sequence and the relative binding positions of each sgRNA.

**Extended Data Fig. 7: Identification of candidate off-target sites by FATE gRNA.**

**a**, Table showing genes, target sequences, genomic locations, mismatch numbers (0, 1, or 2), and A-to-G conversion ratios for candidate sites with high sequence similarity to the FATE gRNA identified by WES in mock- and FATE plasmid–treated groups. **b**, Table showing names, target sequences, genomic locations, mismatch numbers (0 or 1), and A-to-G conversion ratios for candidate sites in non-coding regions, along with the genomic features of each location. **c**, Top : Schematic illustrating the method for identifying the number of overlapping bases with nucleotide substitutions. Bottom : Table showing genomic locations of variants called from WES with ≥200 reads, A-to-G conversion ratios, REF and ALT nucleotide information (A, T, G, C), and sequence similarity to the FATE gRNA in both 5′→3′ and 3′→5′ orientations.

**Extended Data Fig. 8: Integrated characterization of FATE-NMOs by both single-cell profiling and molecular features.**

**a,** Transmission electron microscopy images of day 70 FATE-NMOs showing two major functional compartments. The muscle region displays advanced sarcomeric organization, including aligned myofibrils with distinct Z-lines (Z) and M-bands (M). The neuronal region reveals neurofilaments (NF), mitochondria (Mi), and synaptic vesicles (SV), indicative of mature synaptic structures. Scale bars are indicated in each panel. **b**, Immunofluorescence images of HGPS, Edit, and FATE NMOs stained with Progerin (green), 53BP1 (red), and DAPI (blue). Scale bar, 25 μm. **c**, Transmission electron microscopy (TEM) images showing the nuclear heterochromatin structure in the neuronal region of neuromuscular organoids (NMOs) derived from two isogenic HGPS hPSC pairs, including FATE-NMOs. **d**, Mapping of single-cell transcriptomic data from FATE-treated neural-muscular organoids. **e**, Relative proportions of sub-populations within FATE skeletal muscle cells, visualized in shades of green. **f**, Relative proportions of sub-populations within FATE neural cells, visualized in shades of orange and pink.

**Supplementary Information**

**Supplementary Video 1: 3D confocal imaging of HGPS-NMOs without X-ray irradiation**

Three-dimensional confocal imaging shows progerin (green), 53BP1 (red), and lamin A (grey) in HGPS-NMOs without X-ray irradiation.

**Supplementary Video 2: 3D confocal imaging of HGPS-NMOs with X-ray irradiation**

Three-dimensional confocal imaging shows progerin (green), 53BP1 (red), and lamin A (grey) in HGPS-NMOs 1 hour after 5Gy X-ray irradiation.

**Supplementary Video 3: 3D confocal imaging of Edit-NMOs without X-ray irradiation**

Three-dimensional confocal imaging shows progerin (green), 53BP1 (red), and lamin A (grey) in Edit-NMOs without X-ray irradiation.

**Supplementary Video 4: 3D confocal imaging of Edit-NMOs with X-ray irradiation**

Three-dimensional confocal imaging shows progerin (green), 53BP1 (red), and lamin A (grey) in Edit-NMOs 1 hour after 5Gy X-ray irradiation.

**Supplementary Video 5: 3D confocal imaging of FATE-NMOs without X-ray irradiation**

Three-dimensional confocal imaging shows progerin (green), 53BP1 (red), and lamin A (grey) in FATE-NMOs without X-ray irradiation.

**Supplementary Video 6: 3D confocal imaging of FATE-NMOs with X-ray irradiation**

Three-dimensional confocal imaging shows progerin (green), 53BP1 (red), and lamin A (grey) in FATE-NMOs 1 hour after 5Gy X-ray irradiation.

#### Extended Data Fig. 1

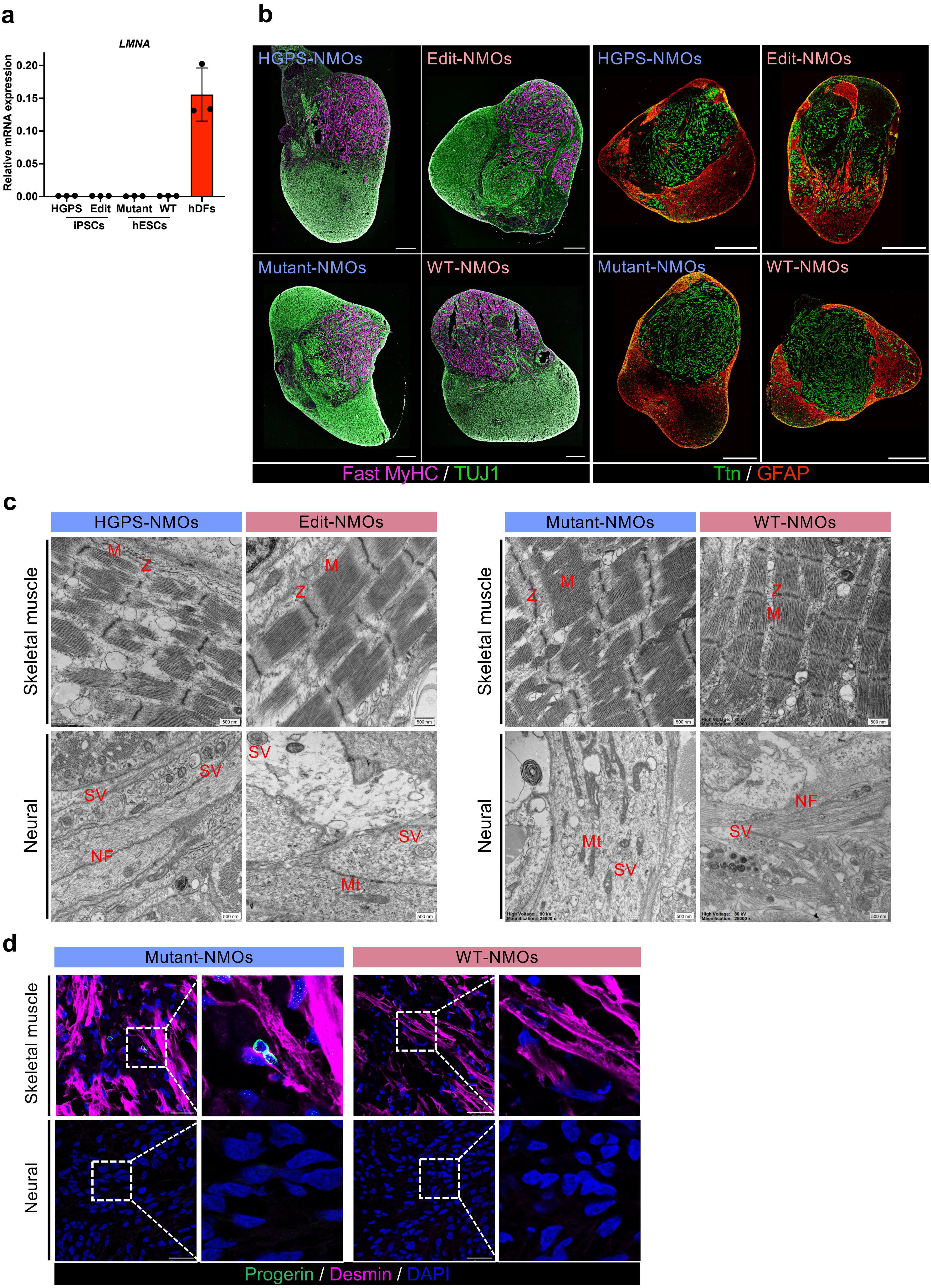

### Extended Data Fig. 2

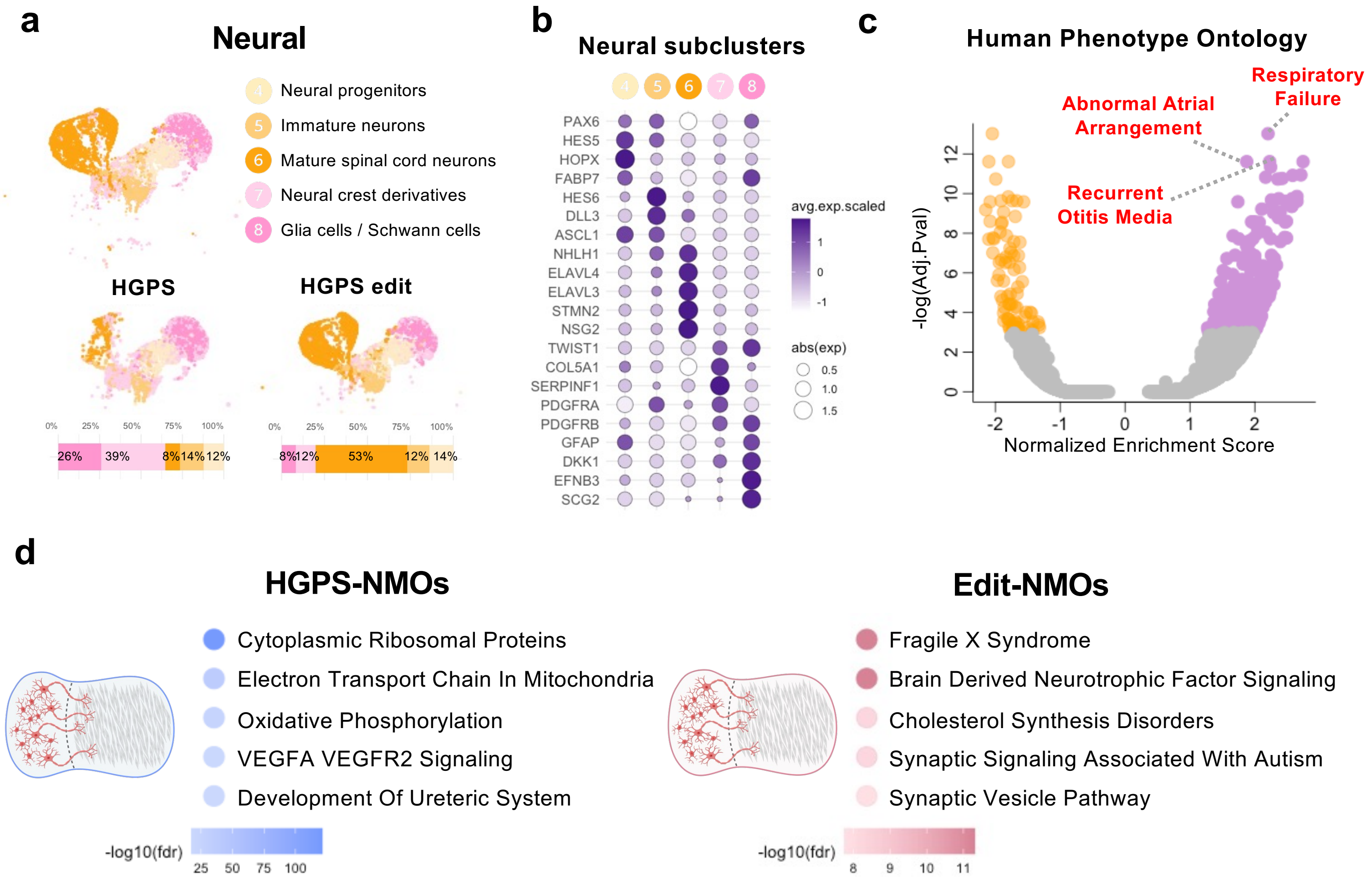

### Extended Data Fig. 3

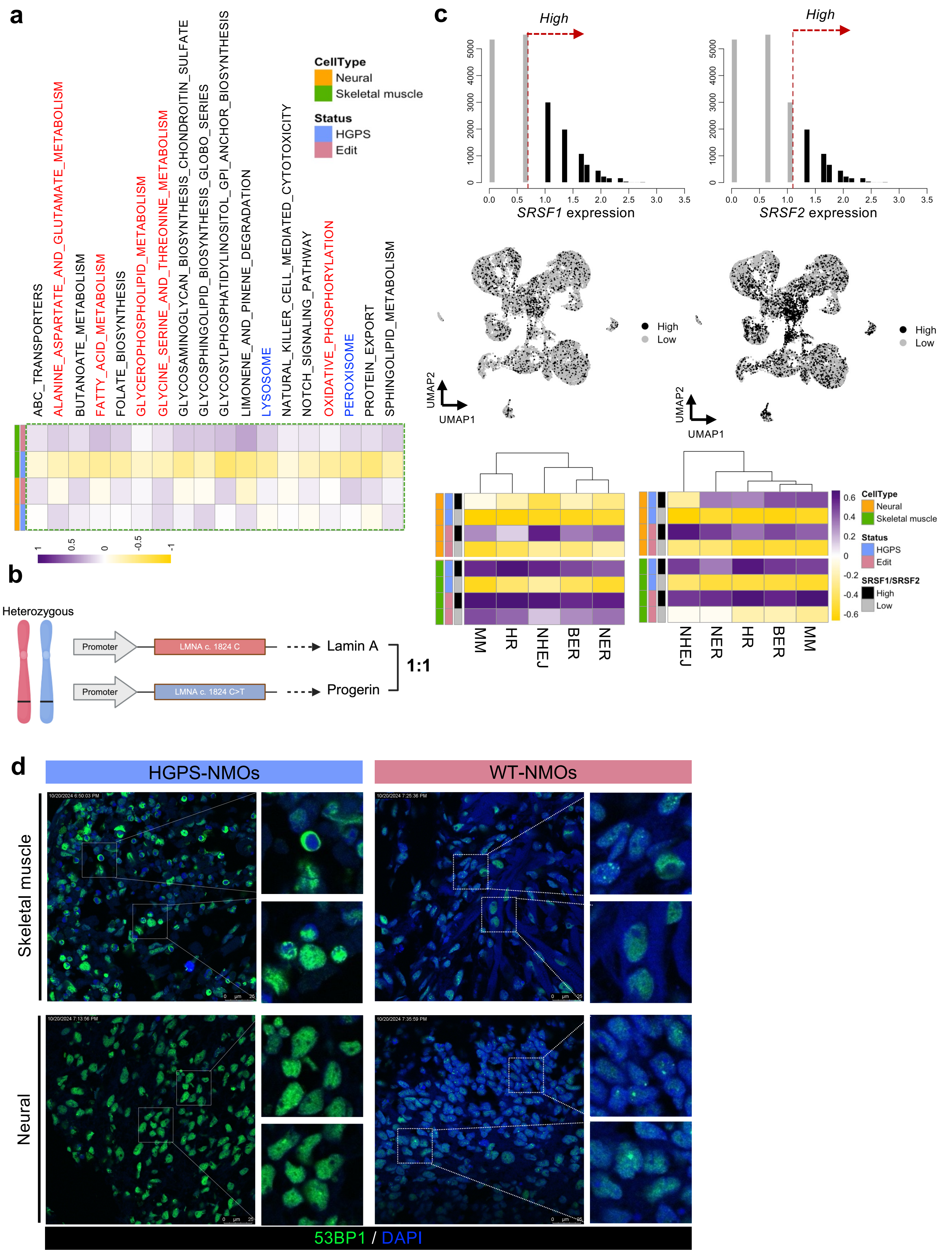

#### Extended Data Fig. 4

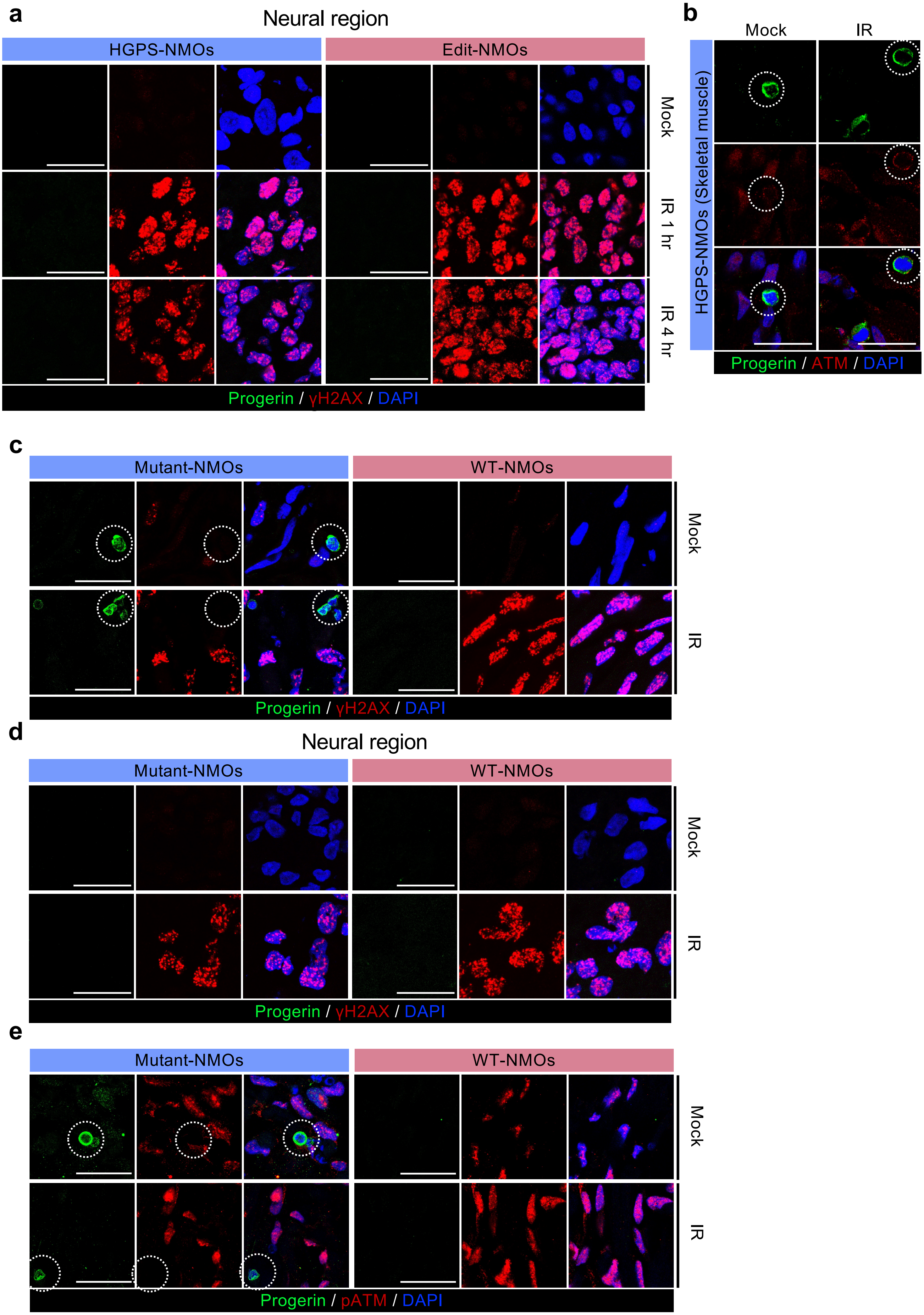

**Extended Data Fig. 5**

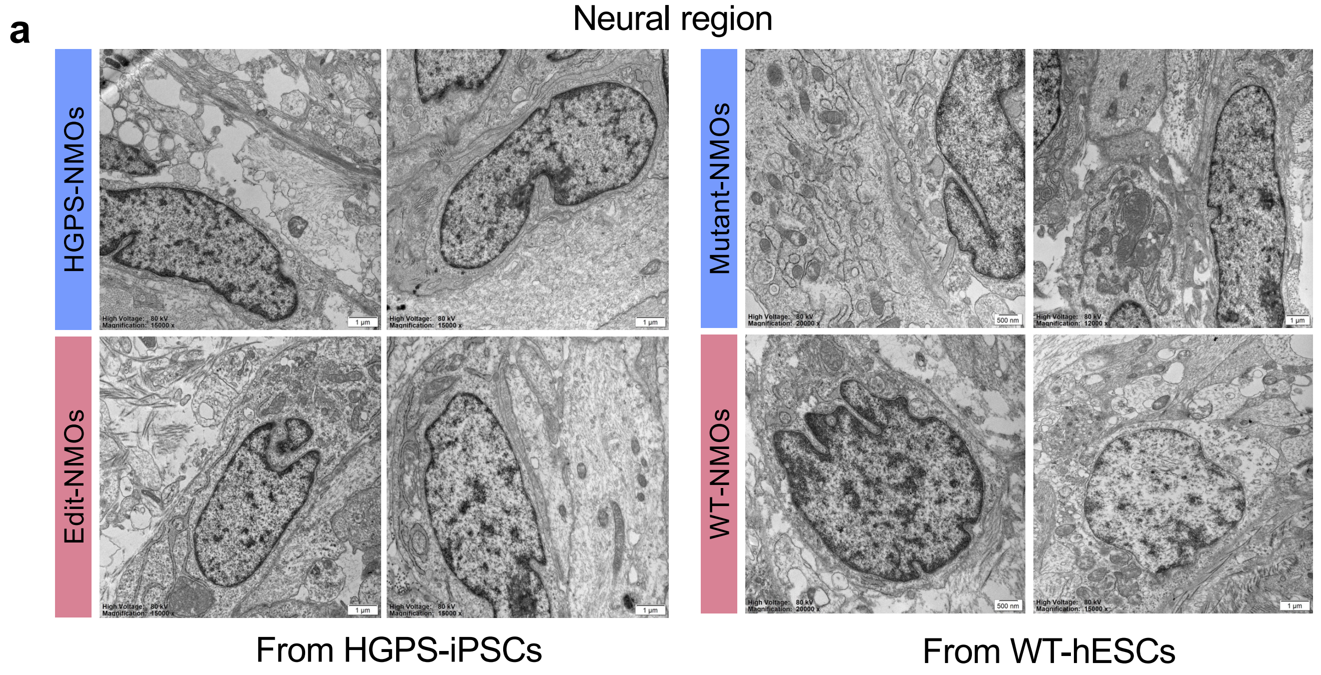

### Extended Data Fig. 6

**a**

| Class | Variant | Protospacer sequence (example) | Editing tool | Concern |
|---|---|---|---|---|
| Classical | c. 1824C>T (antisense G>A) | GGTCCACCCACCTGGGCTCC | ABE | - |
| Non-classical | c. 1821G>A | GTAGGCGGACCCATCTCCTC | ABE | Bystander 'A', Suboptimal |
| | c. 1822G>A | GTGAGCGGACCCATCTCCTC | ABE | Bystander 'A' |
| | c. 1868C>G (antisense G>C) | GCTGCGACTGACCGTGACAC | PE, CGBE | Transversion mutation |
| | c. 1940T>G (antisense A>C) | GCTGGAGTTGCCCAGGCGGT | PE | Transversion mutation |
| | c. 1968G>A | GAACCCAAGTGAGTTGTCTC | ABE | Bystander 'A' |
| | c. 1968+1 G>A | CAGATGAGTTGTCTCTGCTT | ABE | Bystander 'A' |
| | c. 1968+5 G>A | CAGGTGAATTGTCTCTGCTT | ABE | Bystander 'A' |

**b**

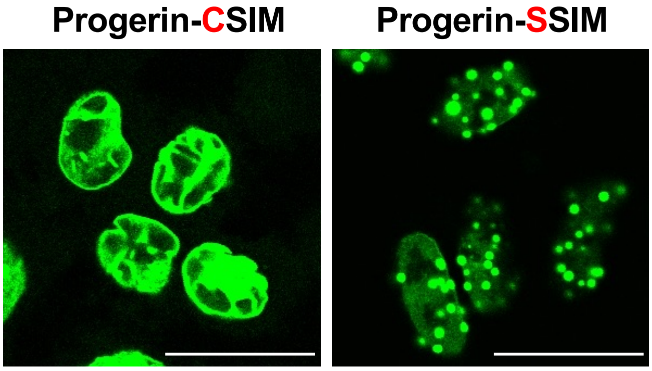


**c**

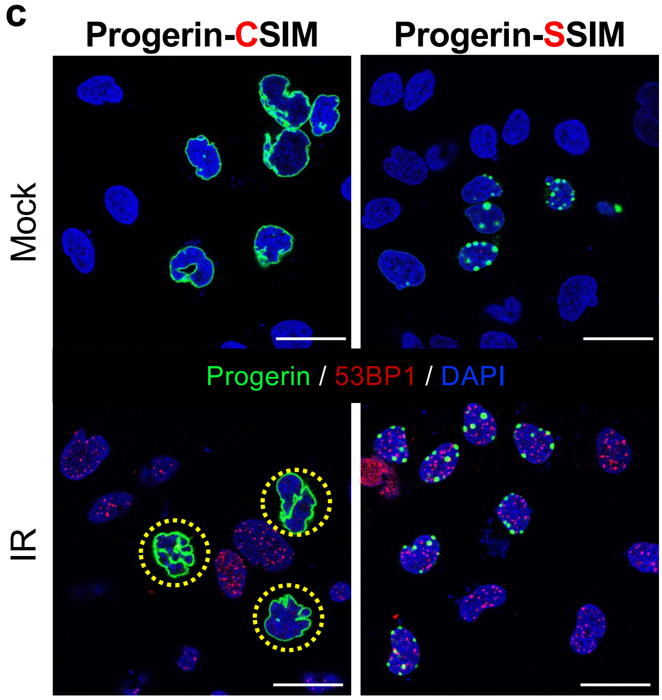


**d**

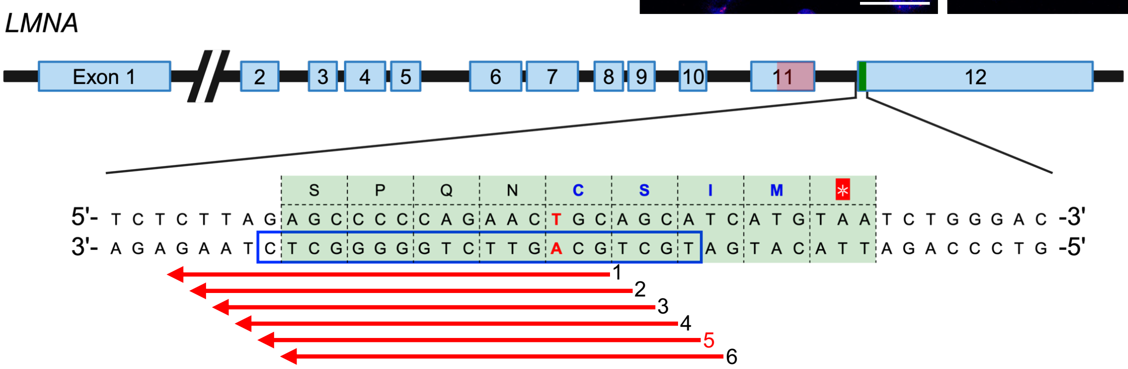


| No | sgRNA Sequence | PAM | ABE version |
|---|---|---|---|
| 1 | GCAGTTCTGGGGGCTCTAAG | AGA | SpRY ABE8e / NG-ABEmax |
| 2 | TGCAGTTCTGGGGGCTCTAA | GAG | SpRY ABE8e |
| 3 | CTGCAGTTCTGGGGGCTCTA | AGA | SpRY ABE8e / NG-ABEmax |
| 4 | GCTGCAGTTCTGGGGGCTCT | AAG | SpRY ABE8e |
| 5 | TGCTGCAGTTCTGGGGGCTC | TAA | SpRY ABE8e |
| 6 | ATGCTGCAGTTCTGGGGGCT | CTA | SpRY ABE8e |

### Extended Data Fig. 7

**a**

| Region | Name | Target | Chromosome | Mismatch | Conversion Ratio |
|---|---|---|---|---|---|
| Exon | *LMNA* | TGCTGCAGTTCTGGGGGCTC | chr1:156139092 | 0 | Mock: P(14):48.3%<br>Treated: P(14):0% |
| Exon | *MYO1E* | TGCTGgAGTTCTGGGtGCTC | chr15:59173819 | 2 | Mock: P(14):0%<br>Treated: P(14):0% |
| Exon | *SLC4A3* | TGCTGgAGTgCTGGGGGCTC | chr2:219629688 | 2 | Mock: P(14):0%<br>Treated: P(14):0% |
| Exon | *TARS2* | TtCTGaAGTTCTGGGGGCTC | chr1:150507161<br>chr1:150507162 | 2 | Mock: P(14):0%<br>Treated: P(14):0%<br>Mock: P(15):0%<br>Treated: P(15):0.22% |
| Exon | *TSKU* | TGCTGCAGTTCTGGaGcCTC | chr11:76797647<br>chr11:76797655 | 2 | Mock: P(6):0%<br>Treated: P(6):0%<br>Mock: P(14):0%<br>Treated: P(14):0% |

**b**

| Region | Name | Target | Chromosome | Mismatch | Conversion Ratio | Note |
|---|---|---|---|---|---|---|
| Exon | LMNA | TGCTGCAGTTCTGGGGGCTC | chr1:156139092 | 0 | Mock: P(14):0.06%<br>Treated: P(14):42.6% | *LMNA*, Positive control |
| Intergenic | Off-Target1 | TGCTGCAGTTCTGGGGGCTC | chr13:98581985 | 0 | Mock: P(14):0.01%<br>Treated: P(14):0.09% | *STK24, SLC15A1* intergenic |
| Intergenic | Off-Target2 | TGCTGCAGgTCTGGGGGCTC | chr12:128780232 | 1 | Mock: P(14):0.01%<br>Treated: P(14):0.01% | *TMEM132C, SLC15A4* intergenic |
| Intergenic | Off-Target3 | TGCTGCAGTTCTGGtGGCTC | chr11:76166363 | 1 | Mock: P(14):0.22%<br>Treated: P(14):18.2% | *UVRAG, WNT11* intergenic |

**c**

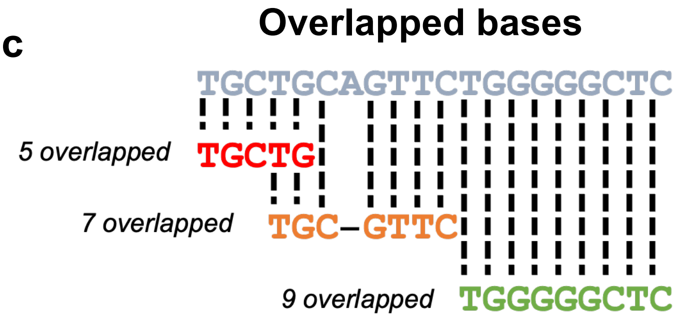


| REF | ALT | DP | Info | Ratio | Ref_values | Alt_values | Overlapped (5′→3′) | Overlapped (3′→5′) | Gene | SNP |
|---|---|---|---|---|---|---|---|---|---|---|
| **WT** | | | | | | | | | | |
| 0 | 421 | 421 | 3;195782558 | 1 | A | G | 5 | 3 | MUC4 | rs28542401 |
| 0 | 642 | 642 | 6;32580950 | 1 | A | G | 5 | 6 | HLA-DRB1 | rs9269768 |
| 361 | 406 | 767 | 13;113816318 | 0.5293351 | A | G | 6 | 5 | TMEM255B | rs67559311 |
| 1 | 230 | 231 | 1;39413694 | 0.995671 | T | C | 7 | 7 | MACF1 | rs604316 |
| 0 | 350 | 350 | 5;711867 | 1 | T | C | 7 | 9 | ZDHHC11B | rs74987061 |
| 451 | 482 | 933 | 5;796162 | 0.5166131 | T | C | 7 | 9 | ZDHHC11 | rs386219 |
| 199 | 289 | 488 | 11;1095004 | 0.5922131 | T | C | 4 | 5 | MUC2 | rs1273413249 |
| 165 | 248 | 413 | 11;1095015 | 0.6004843 | T | C | 4 | 5 | MUC2 | rs1355085893 |
| **Plasmid** | | | | | | | | | | |
| 109 | 132 | 241 | 1;201211020 | 0.5477178 | A | G | 9 | 4 | IGFN1 | rs145517342 |
| 436 | 1009 | 1445 | 3;195783103 | 0.6982699 | A | G | 3 | 4 | MUC4 | rs139925940 |
| 4 | 715 | 719 | 3;195784719 | 0.9944367 | A | G | 5 | 4 | MUC4 | rs3103955 |
| 42 | 1064 | 1106 | 3;195785122 | 0.9620253 | A | G | 4 | 4 | MUC4 | rs3107749 |
| 4 | 510 | 514 | 3;195784654 | 0.9922179 | T | C | 5 | 3 | MUC4 | rs199709753 |
| 196 | 181 | 377 | 1;156139092 | 0.4801061 | T | C | 6 | 20 | LMNA | - |

#### Extended Data Fig. 8

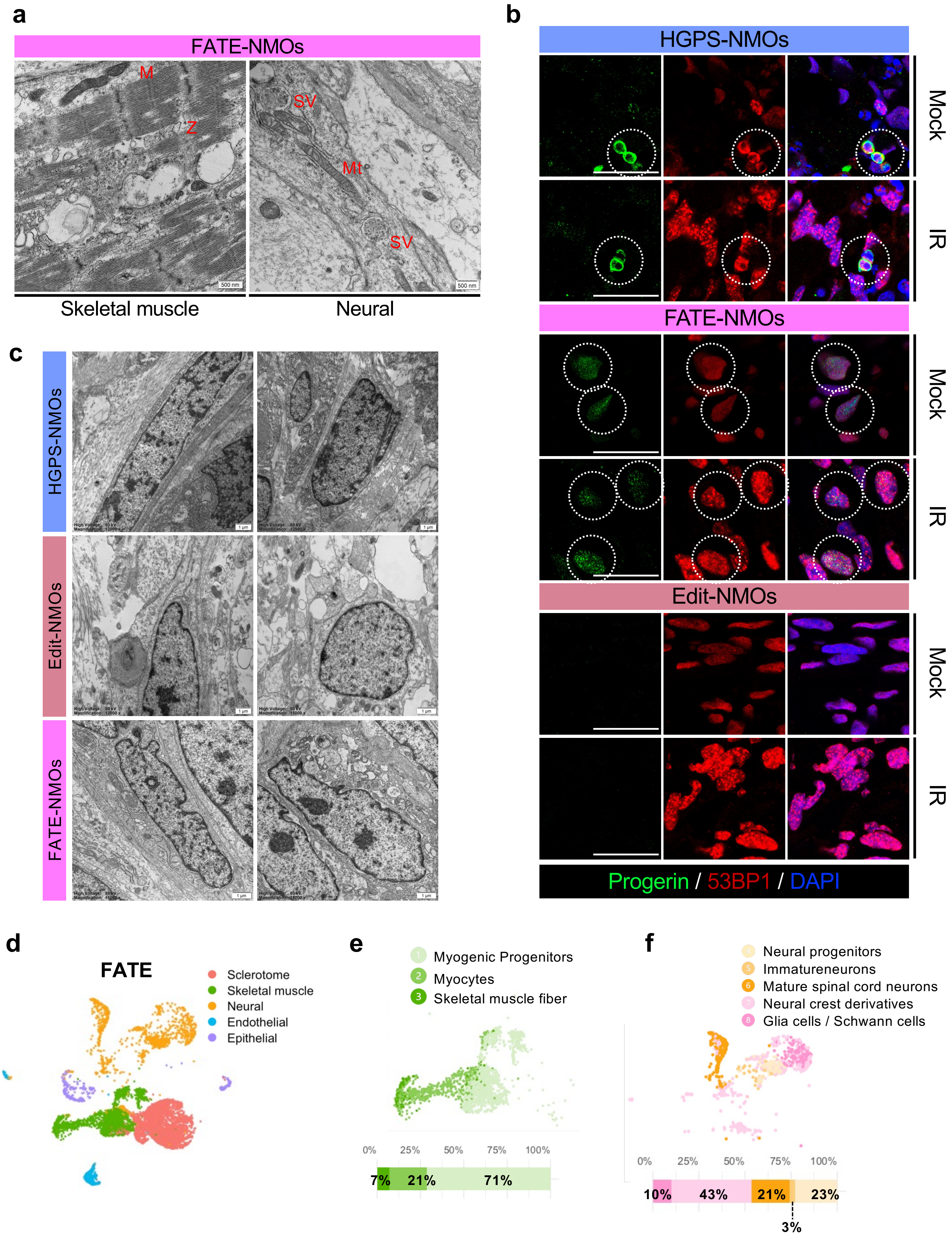
